# A pivotal contribution of proteostasis failure and mitochondrial dysfunction to chromosomal instability-induced microcephaly

**DOI:** 10.1101/2025.03.18.643858

**Authors:** Amanda González-Blanco, Adrián Acuña-Higaki, David Boettger, Marco Milán

## Abstract

Mosaic variegated aneuploidy (MVA), a rare human congenital disorder that causes microcephaly, is characterized by extensive abnormalities in chromosome number and results from mutations in genes involved in accurate mitotic chromosome segregation. To characterize the cellular mechanisms underlying this disease, here we generated a *Drosophila* model of microcephaly caused by the depletion of a single spindle assembly checkpoint (SAC) gene in the neural stem cell (NSC) compartment. We present evidence that loss of stemness - compromised identity and proliferative capacity of NSCs- is the underlying cause of MVA and results in a reduced number of neurons and glial cells. We show that loss of stemness arises from the accumulation over time of an unbalanced number of gains and losses of more than one chromosome, rather than a direct consequence of chromosomal instability-induced DNA damage or the production of simple aneuploidies. We unravel that the negative impact of complex aneuploidies on stemness, a highly energy demanding cellular state, is a result of proteostasis failure and mitochondrial dysfunction. We identify autophagy activation—either directly or through TOR depletion—, overexpression of Radical Oxygen Species scavengers, and restoration of mitochondria proteostasis as genetic interventions capable of dampening the deleterious effects of aneuploidy on NSC identity and brain development.

**Highlights:** SAC depletion in NSCs induces microcephaly through the production of complex aneuploidies Complex aneuploidies compromise the stemness of NSCs

Loss of stemness results from proteostasis failure and mitochondrial dysfunction Boosting proteostasis or mitochondria homeostasis mitigate the negative effects on NSCs

**eTOC Blurb:** Depletion of the spindle assembly checkpoint in the neural stem cell compartment leads to microcephaly. González-Blanco et al. provide evidence that proteostasis failure and mitochondrial dysfunction play a pivotal role in causing microcephaly by compromising the stemness of highly aneuploid neural stem cells.

## Introduction

Mosaic variegated aneuploidy (MVA) is characterized by widespread mosaic aneuploidies, involving different chromosomes and tissues. MVA patients can present growth retardation, early childhood cancer, and microcephaly (García-Castillo et al. 2008; Suijkerbuijk et al. 2010; Pavone et al. 2022; Malumbres and Villarroya-Beltri 2024). MVA is caused by pathogenic mutations in components of the spindle-assembly checkpoint (SAC) and proteins involved in centrosome dynamics during mitosis (Matsuura et al. 2006; Hanks et al. 2004; de Wolf et al. 2021; Langeh et al. 2023; Yost et al. 2017; Carvalhal et al. 2022). In cells from MVA patients carrying bi-allelic mutations in these genes, defects in chromosome segregation cause chromosomal instability (CIN), which results in the generation of aneuploid karyotypes. In mouse models of MVA-driven microcephaly caused by mutations in SAC genes, a dramatic reduction in the neural stem cells (NSCs), their intermediate progenitors, and post-mitotic neurons is associated with massive cell death (Okimoto et al. 2017), which is largely independent of the tumor suppressor gene p53 (Sterling et al. 2023). The *Drosophila* brain, whose development and growth also relies on the proliferative activity of NSCs, has proven to be a valuable model in which to demonstrate a causal relationship between chromosome segregation errors and microcephaly (Gogendeau et al. 2015; Mirkovic et al. 2019; Poulton, Cuningham, and Peifer 2017). By combining changes in centrosome number (amplification or loss) with mutations in SAC genes or by acute targeting of chromosome cohesion, these studies have shown that the resulting brains present extensive abnormalities in chromosome number, a dramatic reduction of the NSC population, and small brain size. Unfortunately, the underlying cellular and molecular mechanisms leading to NSC loss and microcephaly are largely unidentified.

Here we have generated a fly model of microcephaly caused by depletion of a single SAC gene, as occurs in MVA patients. We present evidence that targeted depletion of the SAC in NSCs induces microcephaly in flies and that loss of NSC stemness—compromised identity and proliferative capacity—is the underlying cause of the disease and results in a reduced number of neurons and glial cells. We have characterized the response of the developing brain to CIN-induced DNA damage and different types and levels of aneuploidy and have identified the cellular and molecular mechanisms causing CIN-induced loss of NSC stemness. We present evidence that NSCs are highly resistant to DNA damage and that simple aneuploidies, consisting of single chromosome gains, have a mild impact on the identity and proliferative capacity of these cells and on brain size. In contrast, our results indicate that loss of stemness of SAC-depleted NSCs results from the accumulation, through consecutive cell cycles, of an unbalanced number of gains and losses of more than one chromosome. We performed transcriptomic analysis to unveil a strong effect of CIN on cellular homeostasis. Thus, SAC-depleted brains showed a reduction in genes involved in ribosome biogenesis and RNA metabolism, activation of protein quality control mechanisms (autophagy), and repression of mitochondrial activity. We demonstrate that proteostasis failure—most probably due to aneuploidy-induced gene dosage and proteome imbalance—and mitochondrial dysfunction are major contributors to the deleterious effects of complex aneuploidies on NSC stemness, a highly energy demanding cellular state. Thus, we present evidence that genetic interventions aimed at restoring proteostasis and mitochondrial function can mitigate the negative impact of aneuploidy on NSC biology.

## Results

### Targeted depletion of SAC genes in neural stem cells induces microcephaly

The central region of the *Drosophila* larval brain contains neural stem cells (NSCs), which are of embryonic origin and re-enter the cell cycle after a quiescence period in the first instar larval stage [Figure 1D, (Homem, Repic, and Knoblich 2015)]. Among them, we find the type I neuroblasts (NBs), which divide asymmetrically during the subsequent two larval stages to self-renew and produce ganglion mother cells (GMCs), which will divide once more to give rise to two neurons or glial cells. NBs will differentiate into GMCs or die at the end of larval development, and differentiating neurons and glial cells will build the adult central brain. In humans, MVA-driven microcephaly is caused in half of the patients by bi-allelic mutations in single genes with well-documented roles in spindle assembly checkpoint (SAC) function or correct kinetochore-microtubule attachment (Matsuura et al. 2006; Hanks et al. 2004; de Wolf et al. 2021; Langeh et al. 2023; Yost et al. 2017; Carvalhal et al. 2022). Our first aim was then to address whether targeted depletion of a single SAC gene could also cause a reduction in larval brain size in flies and whether NBs were the most susceptible cells to CIN. We used the *worniu-gal4* driver to induce expression of RNAi forms against the SAC genes *bub3* or *rod* specifically in NBs and to analyze the total number of NBs [labeled by a nuclear GFP/ß-galactosidase fusion protein expressed by the *UAS-GFP-nuc-lacZ* transgene (Shiga, Tanaka-Matakatsu, and Hayashi 1996) and by expression of Deadpan, Dpn], GMCs (labeled by the expression of Prospero, Pros), neurons (labeled by the expression of Elav), and glial cells (labeled by the expression of Repo), as well as the size of the brain lobe (Figure 1A, B, see also Figure S1B). We analyzed control brains and brains subjected to SAC depletion in proliferating NBs at four time points of larval development (48, 62, 72 and 96 h after egg laying, AEL). In control flies not subjected to SAC depletion, NB number in the central brain remained constant throughout larval development, and the number of GMCs and the size of the resulting brain increased as a result of the active proliferative capacity of NBs (Figure 1A, A’). In contrast, in flies subjected to SAC depletion, the number of NBs gradually reduced (Figure 1A, A’). This gradual decrease was also observed upon temporally controlled depletion of the SAC gene using the thermos-sensitive Gal80 system (Figure S1A, A’). Indeed, we noted that the effects on NB number required at least 48 h of SAC depletion (Figure S1A, A’), despite the rapid cell cycles of this population (every 2 h) at these developmental stages (Ito and Hotta 1992). Consistent with this observation, a significant reduction in GMC number and brain size was observed only at later stages of larval development (Figure 1A, A’, see also Figure S1B, B’). As expected, the number of neurons and glial cells in SAC-depleted brains was also reduced when compared to control brains (Figure 1B, B’). These results indicate that the impact of SAC depletion on brain size is a result of a reduced number of GMCs, neurons, and glial cells. To reinforce this notion, we performed lineage tracing of the progeny of control and SAC-depleted NBs using the G-TRACE technique (Evans et al. 2009). As shown in Figure 1C, C’, the area covered by the progeny of SAC-depleted NBs was reduced when compared to control samples.

**Figure 1.**
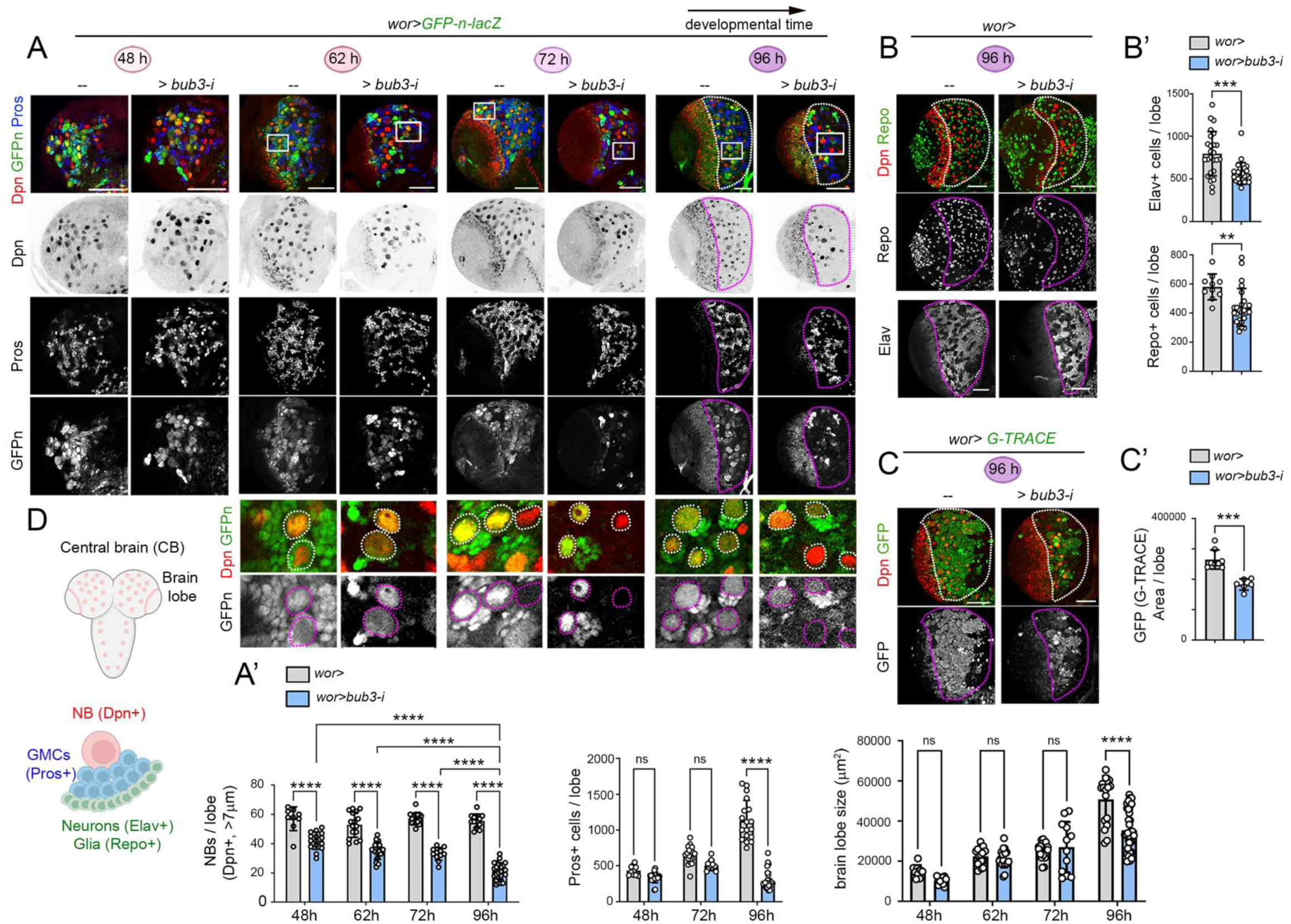
Targeted depletion of SAC genes in neural stem cells induces microcephaly. (**A-C**) Larval brain lobes of larvae of different ages (in hours after egg laying, AEL) expressing an RNAi form for *bub3,* and a nuclear GFP/B-galactosidase fusion protein (**A**) or the G-TRACE cassette (**C**) under the control of the *worniu-gal4* driver, and stained for nuclear GFP (GFP, green and white, **A**, **C**), Dpn (red and black, **A**-**C**), Pros (blue and white, **A**), Repo (green and white, **B**), and Elav (white, **B**). High magnifications of the squared regions are shown in lower panels in **A**, where NBs are labeled by a white line. The central brain region is marked by a dashed line in 96 h AEL brains. Scale bars, 50 µm. (**D**) Drawing of a larval brain and the NB lineage. (**A’**-**C’**) Histograms plotting the number of NBs (**A’**), the number of cells stained for Pros (**A’**), Repo and Elav (**B’**), and GFP (G-TRACE, **C’**) per brain lobe, and the size of the brain lobe (**A’**) of larvae expressing the indicated transgenes and monitored at the indicated ages. Mean and SD are shown. Two-way ANOVA (Šídák’s multiple comparisons test for NBS/lobe and Pros+ cells/lobe, and Tukey’s multiple comparisons test for brain lobe size) (**A’**), or Student’s t-test (**B’, C’**) were performed: * p<0.05, ** p<0.01, *** p<0.001, ****p <0.0001, n.s., not significant. See also Figure S1 and Table S4.

### Targeted depletion of SAC genes induces cell cycle arrest in NSCs

We noticed in control brains expressing the *UAS-GFP-nuc-lacZ* transgene under the control of the *worniu-gal4* driver that the GFP protein was not restricted to NBs and was inherited through perdurance of the Gal4 or GFP proteins by their daughter cells, the GMCs (Figure 1A, lower panels). In contrast, in brains subjected to SAC depletion, the expression of this transgene was very often restricted to NBs and was not observed in the surrounding GMCs, thereby pointing to a failure in the proliferative activity of NBs upon prolonged SAC depletion [see also (Gogendeau et al. 2015) ]. Mitotic activity, monitored with an antibody against a phosphorylated form of histone H3 at serine 10 (PH3) that labels mitotic figures, was first quantified in control and SAC-depleted NBs. As shown in Figure 2A, A’, mitotic activity was compromised in SAC-depleted NBs of mature larval brains. This reduction in mitotic activity was already visible at early larval stages (Figure 2B, B’). We also monitored S phase progression by exposing larval brains to 5-ethynyl-2′-deoxyuridine (EdU) for 45 min and subsequently analyzing its incorporation. The number of EdU-positive NBs was also reduced in SAC-depleted brains when compared to control samples (Figure 2A, A’). Using Fly-Fucci, which relies on fluorochrome-tagged degrons from the Cyclin B (in red) and E2F1 (in green) proteins which are degraded during mitosis or at the onset of the S phase, respectively (Zielke et al. 2014) , we observed that the number of cells not expressing either of these markers and potentially arrested in late G1 were largely increased in SAC-depleted NBs (Figure 2C). A low percentage of NBs also expressed Dacapo (Dap, Figure 2D, D’), the ortholog of p21/p27 in flies that negatively regulates the G1-S transition (de Nooij, Letendre, and Hariharan 1996). However, co-depletion of Dap did not restore the GMC number or brain size (Figure 2D’), thereby pointing to irreversible cell cycle arrest in SAC-depleted NBs. All these data indicate that not only the loss of NBs loss but most probably also their reduced proliferation contribute to CIN-induced microcephaly.

**Figure 2.**
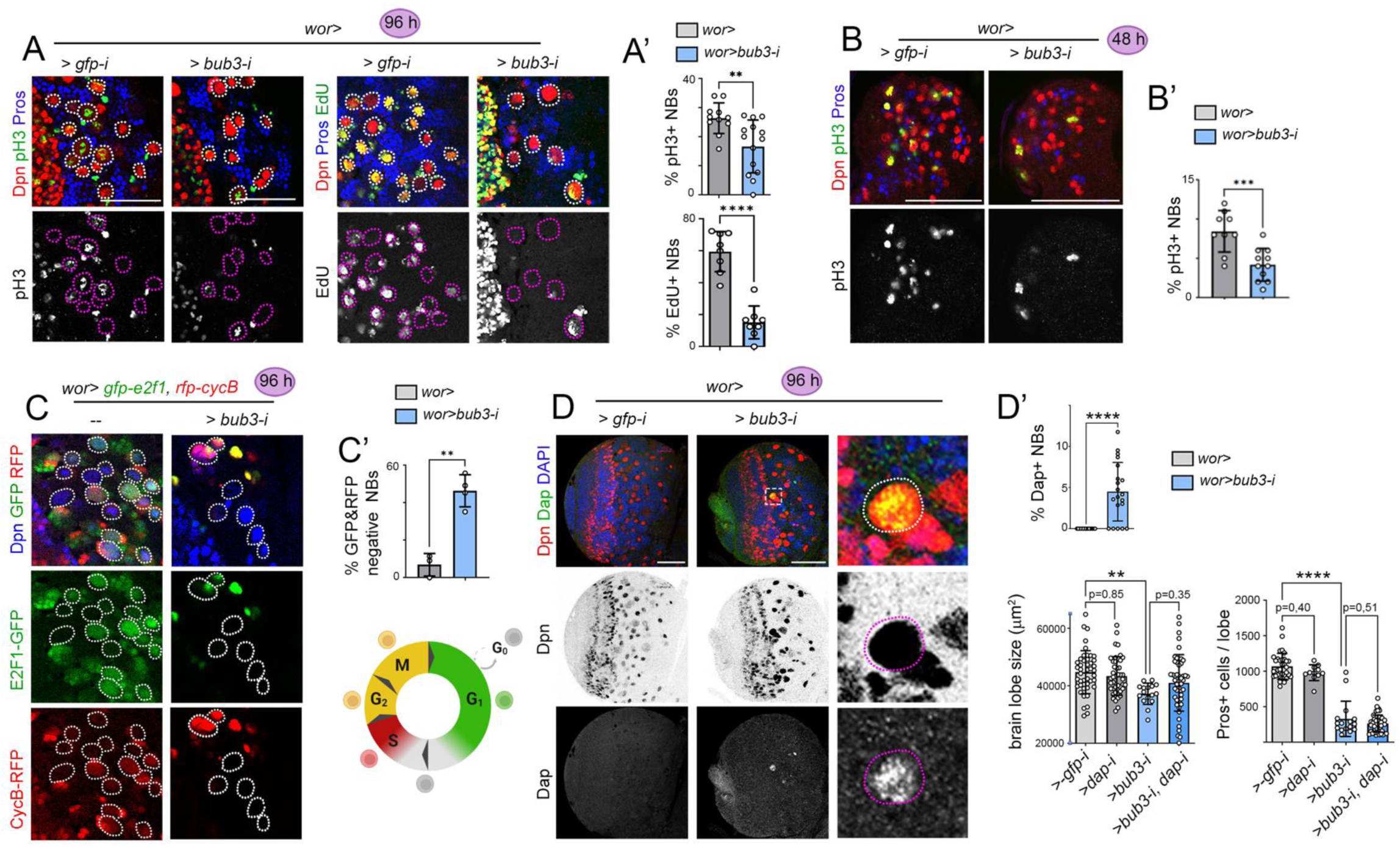
Targeted depletion of SAC genes in neural stem cells induces a cell cycle arrest. (**A-D**) Larval brain lobes exposed to chronic expression of an RNAi form for *gfp* (as controls) or *bub3* under the control of the *worniu-gal4* driver and monitored at 96 h (**A**, **C**, **D**) or 48 h AEL (**B**) to visualize the expression of Dpn (red in **A**, **B**, red or black in **D**, and blue in **C**), Pros (blue, **A**, **B**), pH3 (green and white, **A**, **B**), EdU (green and white, **A**), FlyFucci (GFP-E2F1 in green, RFP-CycB in red, **C**), and Dap (green or white, **D**). **A** and **C** are high magnifications of central brain lobes where NBs are marked by a dashed line. High magnification of the squared region in **D** is shown in the right panel. Scale bars, 50 µm. (**A’**, **B’**, **C’**, **D’**) Histograms plotting the percentage of NBs positive for pH3 or EdU (**A’**, **B’**), negative for GFP-E2F1 and RFP-CycB (**C’**), or positive for Dap (**D’**), and brain lobe size and number of GMCs per lobe (**D’**). Mean and SD are shown. Student’s t-test (**A’, B’, C’**), Mann-Whitney test (for % Dap+ NBs) (**D’**), one-way ANOVA (for brain lobe size and Pros+ cells/lobe, Tukey’s multiple comparisons test) (**D’**) were performed: * p<0.05, ** p<0.01, *** p<0.001, ****p <0.0001, n.s., not significant. See also Table S4.

### Single aneuploidies have a minor effect on NSC number and proliferative capacity

The observation that the effects of SAC depletion on NB number required at least 48 h (Figure 1A, A’ and S1B, B’), despite the rapid cell cycles of this population [every 2 h, (Ito and Hotta 1992) ], pointed to a delayed response of these cells to aneuploidy (Mirkovic et al. 2019). Alternatively, this observation might suggest that NB loss and cycle arrest is a result of a high level of CIN-induced aneuploidy or DNA damage accumulated through consecutive cell cycles. We thus analyzed, in this and the following section, the contribution of different types of aneuploidy and DNA damage to the observed phenotypes. In *Drosophila*, systemic aneuploidies based on trisomies for the X chromosome or the left arm of chromosome II (2L) give rise to mature larvae or pupae that are not able to hatch as adults (Brehme 1937; Fitz-Earle and Holm 1978). Trisomic larvae were generated by crossing females bearing a compound X-chromosome (consisting of two attached X chromosomes that segregate together) or a compound-2L (consisting of two attached 2L arms that segregate together and two free 2R arms) with wild-type males carrying an ubiquitously expressed *GFP* transgene on the X or second chromosomes, respectively, thus labeling trisomic female larvae by the expression of the GFP marker. Trisomies for the X and 2L chromosomes were very stable in larval brains, as shown by the presence of the GFP-expressing chromosome in all cells of the central brain (Figure 3A, B). Given that trisomic larvae underwent developmental delay (data not shown), we quantified the number of NBs and GMCs, as well as brain size throughout the extended larval period and compared these numbers with two types of control euploid larvae, namely those bearing the compound chromosomes and those bearing two unattached chromosomes. The number of NBs was either mildly reduced in the case of X-trisomies (Figure 3A’) or largely unaffected in the case of 2L-trisomies (Figure 3B’). Although NB proliferation (monitored by the number of GMCs at different developmental points) was delayed in trisomic animals when compared to the two controls, the size of the GMC population at the end of the larval period was largely unaffected (Figure 3A, A’, B, B’). A similar trend was observed for brain size (Figure 3A, A’, B, B’). However, brains trisomic for the X chromosome were still smaller than the control brains at the end of the larval period, pointing to the presence of X chromosome-linked triplosensitive loci affecting the growth of other regions of the brain. All these data indicate that trisomies for the X chromosome and 2L arm have a minor impact on NB viability and proliferative capacity, despite the high number of protein-encoding genes that are potentially unbalanced in trisomics for the X (2196 genes) and 2L chromosomes (2657 genes).

**Figure 3.**
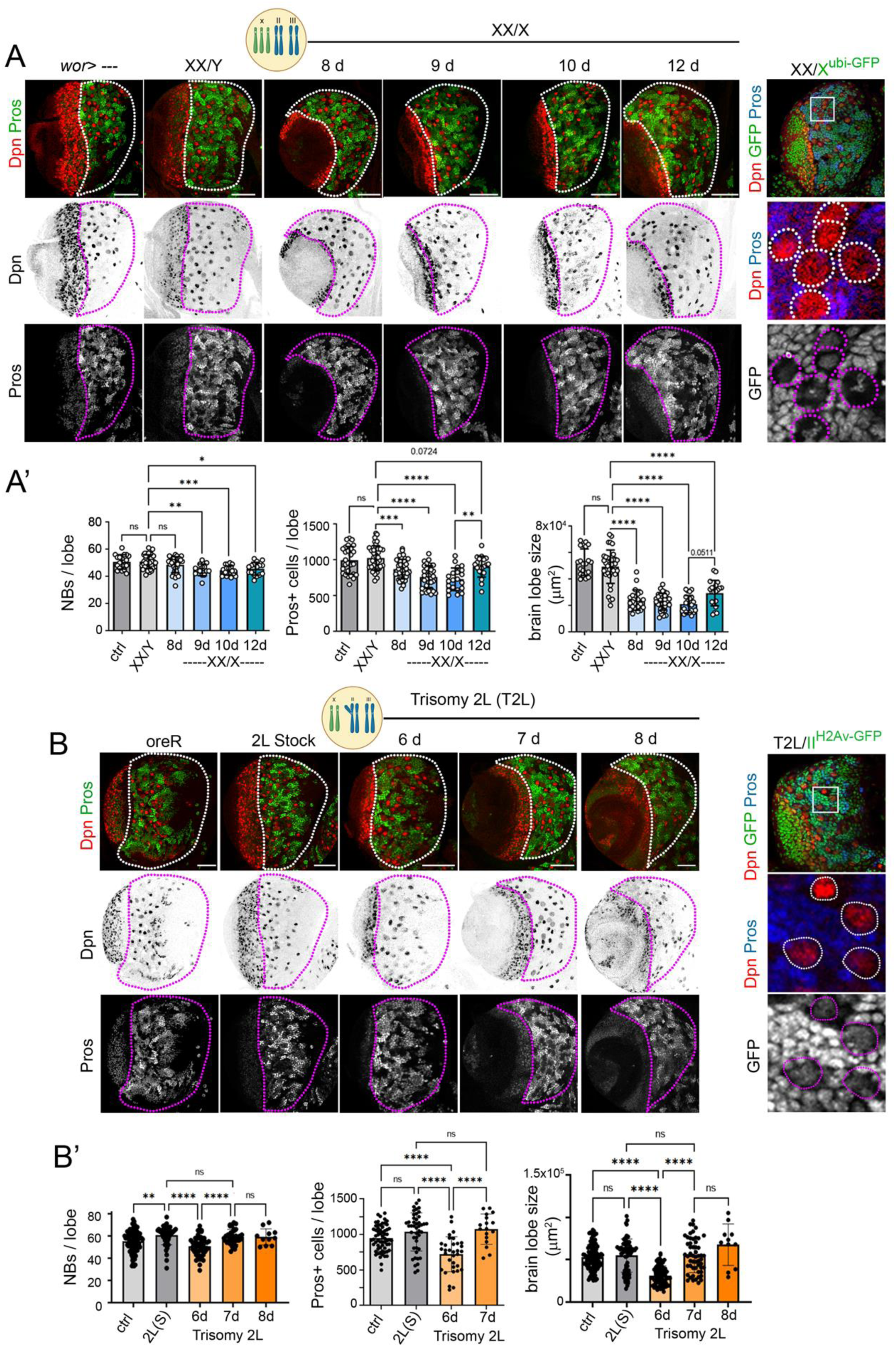
Effects of trisomies on brain development. (**A**, **B**) Larval brain lobes of larvae trisomic for the X (**A**) or 2L (**B**) chromosomes and labeled to visualize the expression of Dpn (red or black), Pros (green, blue or white) and GFP (*ubi-GFP* transgene is located on the X, **A**, or second, **B**, chromosomes). Two controls were used: *wild-type* larvae, and larvae bearing compound chromosomes but maintaining euploidy. The central brain region is marked by a dashed line, and NBs are marked by a dashed line in the high magnification of the squared regions. Scale bars, 50 µm. (**A’**, **B’**) Histograms plotting the number of NBs and Pros-positive cells per lobe, and the size of the brain lobe of larvae of the indicated genotypes and monitored at the indicated ages (in days AEL). Mean and SD are shown. One-way ANOVA (**A’**, **B’**, Tukey’s multiple comparisons test) was performed: * p<0.05, ** p<0.01, *** p<0.001, ****p <0.0001, n.s., not significant. See also Figure S2 and Table S4.

The presence of a single X chromosome in *Drosophila* males causes an imbalance between X-linked and autosomal genes, which is solved by the dosage compensation mechanism (DCM, Figure S2A). The Male-Specific Lethal complex (MSLc), which can be visualized in male tissues by the expression of the Msl2 protein (Figure S2B), targets the X chromosome in males, leading to de-compaction of the chromatin fiber and increased gene transcription to a similar level to that observed in autosomal chromosomes. In females, RNA binding protein Sex lethal (Sxl) induces the translational repression of Msl2 [Figure S2A, B, (Lucchesi and Kuroda 2015)]. Experimental depletion of Msl2 and Sxl in male and female tissues, respectively, causes an X chromosome-wide gene expression imbalance with respect to autosomes [Figure S2C, (Hamada et al. 2005; Alekseyenko et al. 2012)]. In contrast to the deleterious effects of these manipulations on the growth and development of epithelial tissues in *Drosophila* (Clemente-Ruiz et al. 2016), the number of NBs was largely unaffected upon Msl2 depletion in males or Sxl depletion in females (Figure S2D). These results reinforce the conclusion regarding the minor impact of single trisomies and their resulting gene dosage imbalance on NB viability and extend this conclusion to single monosomies (in terms of gene expression) caused by Msl2 depletion in male brains.

### DNA damage has a delayed impact on NSC number and proliferative capacity

Lagging chromosomes are frequent in cells with SAC defects and are likely to be damaged (Vitre and Cleveland 2012). Consequently, the DNA damage response (DDR) pathway can be activated (Janssen et al. 2011). Interestingly, SAC-depleted NBs showed increased levels of phosphorylated H2Av [pH2Av, a target of ATM and ATR in flies (Dekanty et al. 2015)], and activation of p53 [monitored by a GFP activity sensor [(Zhang et al. 2014), Figure 4A)]. However, depletion of DDR genes [*Dp53*, *ATR* (*mei41* in flies) or *chk2* (*grapes* in flies) by RNAi or the use of a dominant negative form of Dp53 (eg. Dp53^H159N^) did not have a major impact on the number of SAC-depleted NBs, and gave rise only to a modest reduction in the size of CIN brains (Figure 4B, B’). Depletion of DDR genes alone did not cause any major effect on brain size or NB number (Figure 4B, B’). To further address the contribution of DNA damage to the observed CIN-induced phenotype, we examined whether DNA damage caused by high doses of ionizing radiation (X-ray, IR, 4500 rads) would phenocopy the CIN-induced reduction in the number of NBs or their progeny, as well as the decrease in brain size. Despite strong activation of the DDR pathway, monitored by increased levels of pH2Av in larval brains 5 h after IR, cell death (monitored with an antibody against the cleaved form of the effector Caspase Dcp1, cDcp1), NB number, and mitotic activity were largely unaffected when compared to control non-irradiated brains (Figure 4C, C’). In contrast, epithelial cells of the larval wing primordium subjected to a similar dose of IR showed high levels of cDcp1 and low mitotic activity 5 h after exposure (Figure S3A-C). cDcp1 and mitotic activity in the wing primordium were restored to control levels 3 days after IR exposure (Figure S3A-C). However, the number of NBs and GMCs was clearly reduced 3 days after IR exposure, at a time at which pH2Av levels were reduced to those of control brains [Figure 4C, C’, D, D’, see also (Wagle and Song 2020)]. The temporal delay observed between H2Av phosphorylation, which marks the presence of double-strand breaks (DSBs), and loss of NBs and reduction in GMC number suggests that these phenotypes are an indirect consequence of the presence of DSBs in the tissue, whereby unrepaired DNA would cause genome rearrangements and aneuploidy. Fluorescence in situ hybridization (FISH) analysis of NBs 3 days after IR to count the number of copies of chromosome II revealed a clear increase in the percentage of NBs losing or gaining at least one chromosome (Figure 4E, E’, note that fly chromosomes in interphase are closely attached in normal conditions, thus the presence of a single chromosome in control NBs). These data suggest that the delayed impact of DNA damage to the brain phenotype relies on the production of aneuploid karyotypes, and they point to a role of DNA damage in enhancing aneuploidy levels in CIN brains rather than an immediate effect on the viability or proliferative capacity of NBs.

**Figure 4.**
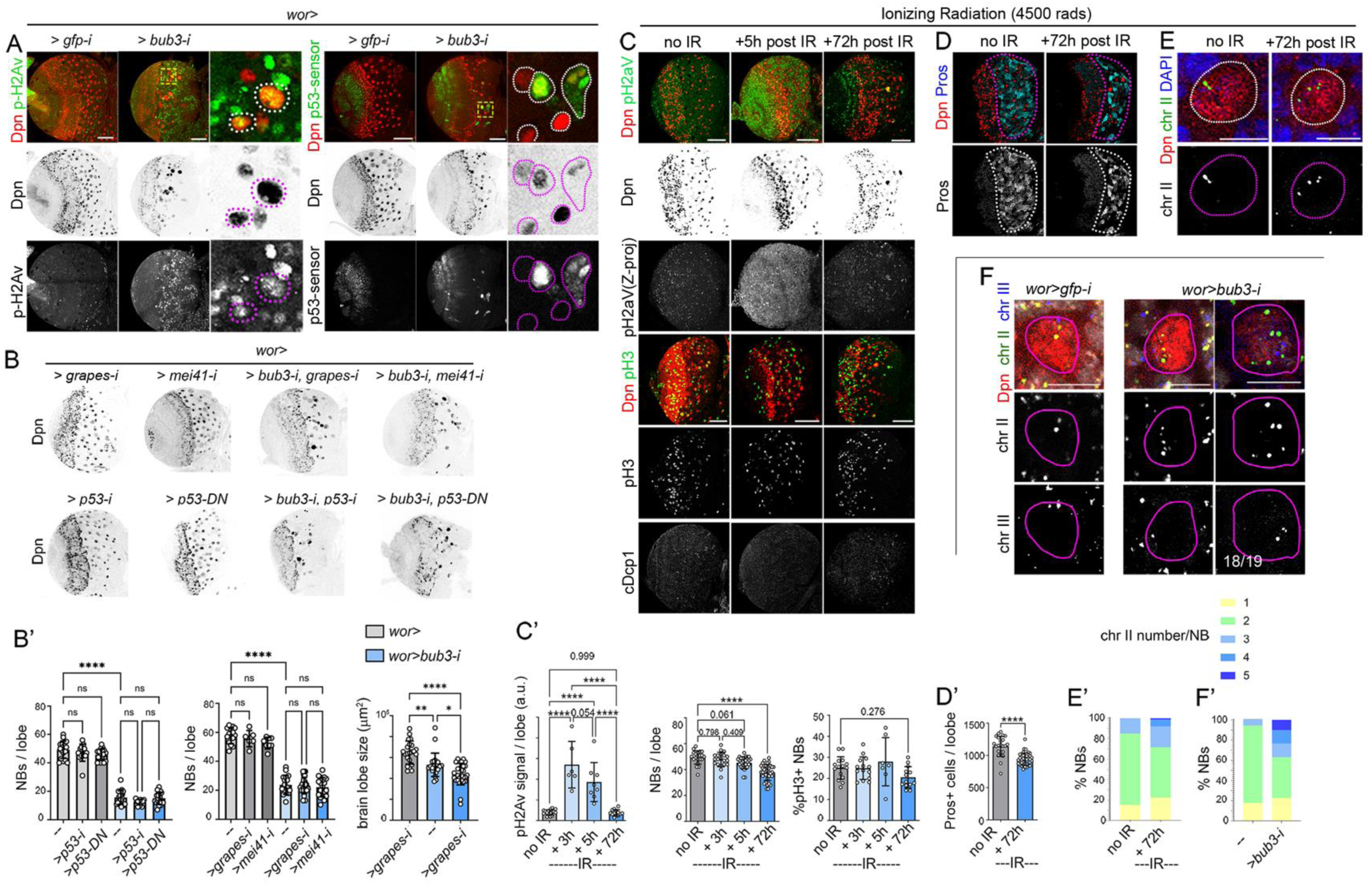
A discrete role of DNA damage on neural stem cell viability. (**A**-**F**) Larval brain lobes expressing the indicated transgenes under the control of the *worniu-gal4* driver (**A**, **B**, **F**) or subjected to ionizing radiation 5 or 72 h before dissection (**C**-**E**) and labeled to visualize the expression of Dpn (red or black, **A**-**F**), pH2Av (green or white, **A**, **C**), p53 sensor (green or white, **A**), pH3 (green or white, **C**), cDcp1 (white, **C**), Pros (blue or white, **D**), chromosome II (green and white, **E**, **F**), and chromosome III (blue and white, **F**). NBs are marked by a dashed line in the high magnification of the squared regions in **A** and in **E**, and in **F**. Central brain is marked by a dashed line in **D**. The ratio of NBs with an unbalanced number of chromosomes is shown in **F**. Scale bars, 50 µm. (**B’**-**F’**) Histograms plotting the number of NBs (**B’**, **C’**), Pros-positive cells (**D’**) and pH2Av signal (**C’**) per brain lobe, brain lobe size (**B’**), and percentage of pH3-positive NBs (**C’**) or NBs containing different numbers of chromosome II (**E’**, **F’**) of larvae subjected to transgene expression or ionizing radiation. Mean and SD are shown. One-way ANOVA (**B’, C’**, Tukey’s multiple comparisons test), Student’s t-test (**D’**), and two-way ANOVA (E’, F’, Šídák’s multiple comparisons test) were performed: * p<0.05, ** p<0.01, *** p<0.001, **** p<0.0001, n.s., not significant. See also Figure S3 and Table S4.

Our experimental data on the minor effects of simple aneuploidies on NB viability and proliferative capacity (Figure 3 and Figure S4) and on the proposed effects of DNA damage in enhancing the aneuploidy levels of CIN brains (Figure 4) point to complex aneuploidies, consisting of multiple losses and gains of chromosomes, as those responsible for NB loss, cell cycle arrest, and CIN-induced microcephaly. Consistently, FISH analysis of NBs to count the number of copies of chromosomes II upon chronic induction of CIN revealed an increase in the percentage of NBs losing or gaining at least one chromosome (Figure 4F, F’). Moreover, by using two distinct FISH probes to label chromosomes II and III, we observed that most NBs (18/19) accumulated more than one chromosome and showed an imbalanced number of chromosomes. This finding points to this imbalance as the major contributor of CIN-induced microcephaly (Figure 4F). These results indicate that complex aneuploidies accumulated through consecutive cell cycles and consisting of multiple gains and losses of chromosomes exert a negative impact on NB viability and proliferative capacity.

### CIN brains show transcriptional changes in protein quality control mechanisms and translation

To identify the molecular mechanisms underlying the negative impact of complex aneuploidies on NB survival and proliferation, we characterized the transcriptional profile of brains subjected to chronic depletion of the SAC gene *bub3* and compared it with that of control brains. Among the most upregulated genes (>1.5 fold, 28 genes, Table S1) in CIN brains, we observed the presence of four members of the Heat Shock Protein 70 (Hsp70) superfamily of chaperones (*hsp70Ba*, *hsp70Bb*, *hsp70Bc*, and *hsp70Bbb*, Figure 5A), mediating protein folding, and Glutathione S transferase E8, which belongs to a protein family involved in ROS scavenging. Gene set enrichment analysis also pinpointed genes involved in vesicle trafficking, autophagy, mitophagy, and protein folding as classes of genes whose expression profiles were also significantly upregulated in CIN brains (Figure 5C, D, Figure S4B and Tables S2 and S3). Among the most downregulated genes (>1.5 fold, 33 genes, Table S1) in CIN brains, we found genes encoding transcription factors involved in neuronal or glial specification, including, as expected, *prospero* [e.g., *castor*, *datilografo*, *huckebein*, *bric a brac 1* ), genes involved in mitochondrial homeostasis [e.g., *dj-1beta*, (Meulener et al. 2006; De Lazzari et al. 2023)] and *bub3* (Figure 5A and Table S1). Gene set enrichment analysis also highlighted genes involved in RNA metabolism, nucleolar function, and mitochondrial ribosomes as classes whose expression profiles were also significantly downregulated in CIN brains (Figure 5C, D, Figure S4B, and Tables S2 and S3). Interestingly, the overall transcriptional profile of the mitochondrial genome was also reduced in these brains (Figure 5B). All these results point to a reduction in translation (e.g., ribosome biogenesis) and RNA metabolism, activation of protein quality control mechanisms (autophagy), and repression of mitochondria activity as the major responses of developing brains to CIN-induced aneuploidy. We next validated some of these responses by immunochemistry or the use of activity reporters and characterized their functional contribution to the deleterious effects of complex aneuploidies on NBs.

**Figure 5.**
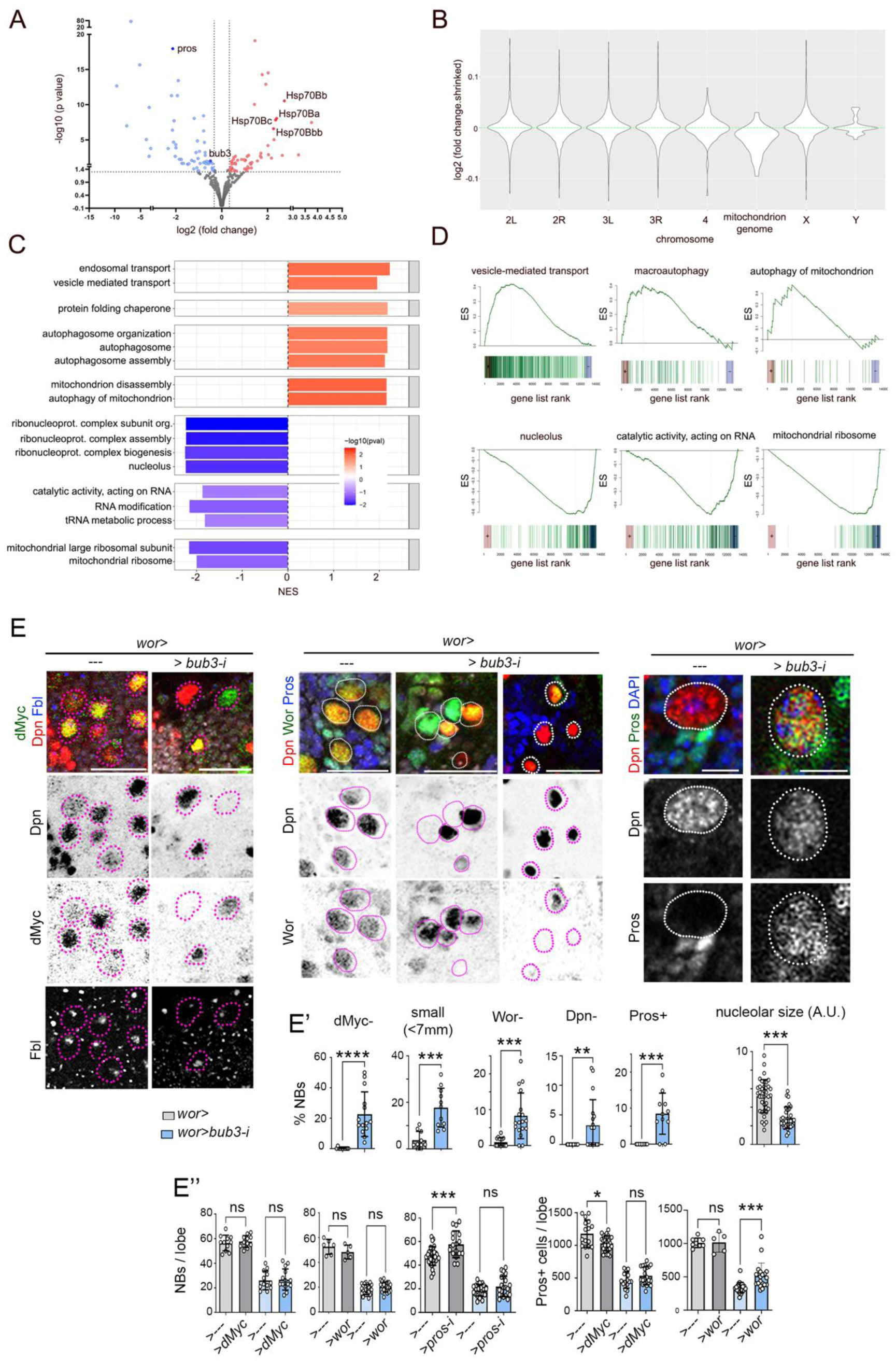
Complex aneuploidies induce loss of stemness. (**A**-**D**) Transcriptional changes of larval brain lobes exposed to chronic expression of an RNAi form of *bub3* (SAC depletion) when compared to controls (expressing *gfp-RNAi*) under the control of the *worniu-gal4* driver and monitored at 96 h AEL. In the volcano plot (**A**), significantly up-(red) or down-regulated (blue) genes (P < 0.05, abs (fold change) ≥ 1.25) are highlighted. Relevant genes are labeled with their corresponding symbols. Fold changes and p-values were calculated using the DESeq2 R package. Violin plot in (**B**) represents the fold change of those genes located in each of the chromosomes and the mitochondrial genome. Bar plot in (**C**) represents GO and KEGG pathway enrichment analysis for up- (red) and down-regulated (blue) gene sets. Statistics from the roastgsa enrichment algorithm of selected pathways are shown. P-value sign corresponds to that of the normalized enrichment score (NES). See the methods section for details on methodology. Gene set enrichment analysis of some GO and KEGG pathways are shown in **D**. (**E**) High magnifications of larval brain lobes expressing the indicated transgenes under the control of the *worniu-gal4* driver and labeled to visualize the expression of Deadpan (Dpn, red or black), dMyc (green or black), and Fibrilarin (Fbl, blue or white). NBs are marked by a dashed line. Scale bars, 25 µm (1^st^ and 2^nd^ panel) and 7 µm (3^rd^ panel). **(E’)** Histograms plotting the percentage of NBs losing dMyc expression, percentage of cells <7 µm Dpn+, percentage of NBs losing Wor expression, percentage of NBs losing Dpn expression, percentage of NBs (Dpn+, >7 µm) with Pros expression, and size of nucleolus according to Fbl signal. The number of NBs and Prospero-positive cells per brain lobe (**E’’**) of larvae subjected to the expression of the indicated transgenes. Mean and SD are shown. Mann-Whitney test (**E**, dMyc-, Dpn-, Pros+), Student’s t-test (**E**, small, Wor-, nucleolar size), and one-way ANOVA (**E’,** Tukey’s multiple comparisons test) were performed:* p<0.05, ** p<0.01, *** p<0.001, **** p<0.0001, n.s., not significant. See also Figure S4 and Table S1-S4.

### Aneuploidy induces loss of stemness in NSCs

In developing NBs, ribosome biogenesis in the nucleolus relies on the activity of the dMyc proto-oncogene, which contributes to their enlarged cell and nuclear size (Rust et al. 2018). Consistent with the observed transcriptional reduction in nucleolar protein-encoding genes in CIN brains (Figure 5C, D), a large proportion of aneuploid NBs lost dMyc expression, showed reduced levels of Fibrilarin [a nucleolar protein involved in RNA processing and regulated by dMyc, (Coller et al. 2000; Orian et al. 2003)], and were small (Figure 5E, E’). dMyc also plays an important role in conferring NB identity by maintaining cortical polarity, a requirement for the intrinsic capacity of NBs to divide asymmetrically and avoid the nuclear entry of the transcription factor Prospero (Rust et al. 2018). Consistently, a proportion of aneuploid NBs showed nuclear localization of Prospero [Figure 5E, E’, see also (Gogendeau et al. 2015; Mirkovic et al. 2019)], and, most interestingly, these NBs also lost the expression of Dpn or Worniu (Figure 5E, E’), two transcription factors involved in promoting NB self-renewal through asymmetric cell division and avoiding premature differentiation (Lai et al. 2012; San-Juán and Baonza 2011). All these observations indicate that the aneuploidy-induced reduction in NB number and proliferative capacity is most probably a result of loss of stemness. Overexpression of dMyc or Worniu did not block the elimination of aneuploid NBs (Figure 5E’’). Only Worniu, when overexpressed, was able to partially rescue the number of Prospero-positive cells in CIN brains (Figure 5E’’), most probably through its role in driving NB cell cycle progression (Lai et al. 2012). Neither did depletion of the pro-differentiation transcription factor Prospero (Pros) rescue the reduction in the number of aneuploid NBs caused by CIN (Figure 5E’’). Apoptosis has alternatively been proposed to play a role in the reduction in brain size caused by changes in centrosome number and SAC depletion (Poulton, Cuningham, and Peifer 2017). To quantify the levels of apoptosis in NBs subjected to *bub3* depletion, we monitored activation of the apoptotic machinery with an antibody that detects the cleaved form of the effector Caspase Dcp1 (cDcp1) and an activity reporter of effector Caspases Dcp1 and Drice that becomes fluorescent upon caspase-driven cleavage [GC3Ai, (Schott et al. 2017)]. Whereas the percentage of control NBs positive for either of these two markers remained close to zero throughout larval development, a low but significant percentage of NBs subjected to CIN were positively labeled by these markers (Figure S4C, C’’). Inhibition of apoptosis with the baculovirus protein p35, an inhibitor of effector caspases in *Drosophila*, restored the number of NBs subjected to CIN only at early stages of development. However, the dramatic reduction in NB number in mature brains was still observed [Figure S4C’’, see also (Gogendeau et al. 2015)]. In contrast, the reduction in the number of Prospero-positive cells was partially restored in mature brains subjected to CIN and expressing p35 (Figure S4C’’). The capacity of p35 to partially rescue the number of Prospero-positive cells can be explained by the perdurance of the protein that is inherited by progenitor cells and points to a role of apoptosis in eliminating progenitor cells that become aneuploid. All these results indicate that the aneuploidy-induced loss of NB identity and proliferative capacity is a consequence of loss of stemness but not cell death or premature differentiation, as previously proposed (Gogendeau et al. 2015; Poulton, Cuningham, and Peifer 2017). Consistent with this notion, *prospero* depletion combined with apoptosis inhibition did not rescue the dramatic reduction in NB number in mature brains subjected to CIN either (Figure S4D, D’). We then addressed whether the loss of stemness - a highly energy demanding cellular state - observed in highly aneuploid NSCs is caused by proteostasis failure and mitochondrial dysfunction.

### Loss of stemness is a consequence of proteostasis failure and mitochondrial dysfunction

As a follow-up of the identification of a general upregulation of genes involved in protein folding, vesicle trafficking and autophagy in CIN brains, by mean of RNAseq, we next monitored the activity of the two main protein quality control mechanisms, namely the ubiquitin-proteasome system (UPS) and autophagy, in aneuploid NBs. To characterize the *in vivo* activity of the proteasome in these cells, we used CL1-GFP, a fusion protein created by attaching a proteasome degradation signal to GFP (Pandey et al. 2007). Whereas CL1-GFP was rapidly degraded in control NBs and no accumulation was observed, this protein accumulated in aneuploid NBs (Figure 6A), pointing to a situation of near saturation of the proteasome in these cells. To reinforce this notion, we used Htt25Q, a fusion protein consisting of human Huntingtin with a polyQ stretch containing only 25 glutamines and fused to a Cerulean fluorescent protein (Ramdzan et al. 2017). Htt25Q does not form aggregates nor does it cause toxicity on its own. However, the presence of protein aggregates in the cell can seed Htt25Q polymerization (Serpionov et al. 2016). Whereas Htt25Q showed diffuse expression in control NBs (Figure 6B) and did not alter the number of NBs (Figure 6E and Figure S5B), it accumulated in large foci in aneuploid NBs (Figure 6B) and caused a further reduction in the size of this population (Figure 6E and Figure S5B). Together with the UPS, autophagy—a pathway responsible for the lysosomal degradation of cellular components—plays an essential role in maintaining protein homeostasis (proteostasis). High levels of autophagy were induced in aneuploid NBs, as visualized by the punctated accumulation of ChAtg8a [a mCherry-tagged form of Atg8a, which becomes membrane-associated under autophagy induction and localizes to the site of autophagosome nucleation (Hegedűs et al. 2016)] in these cells (Figure 6C). In control brains, small Atg8a-positive puncta were observed only in small cells around the NBs but never in the NBs themselves (Figure 6C, arrowheads). Interestingly, boosting autophagy (by overexpressing Atg1 or depleting the activity of Tor through Rheb depletion or the use of Tor mutants in heterozygosis) led to significant rescue of the number of aneuploid NBs in mature CIN brains, without causing any major effect in otherwise control brains (Figure 6E and Figure S5A). These results indicate that proteostasis failure, most probably due to an imbalance in the proteome caused by aneuploidy, functionally contributes to the reduction in the number of NBs in CIN brains.

**Figure 6.**
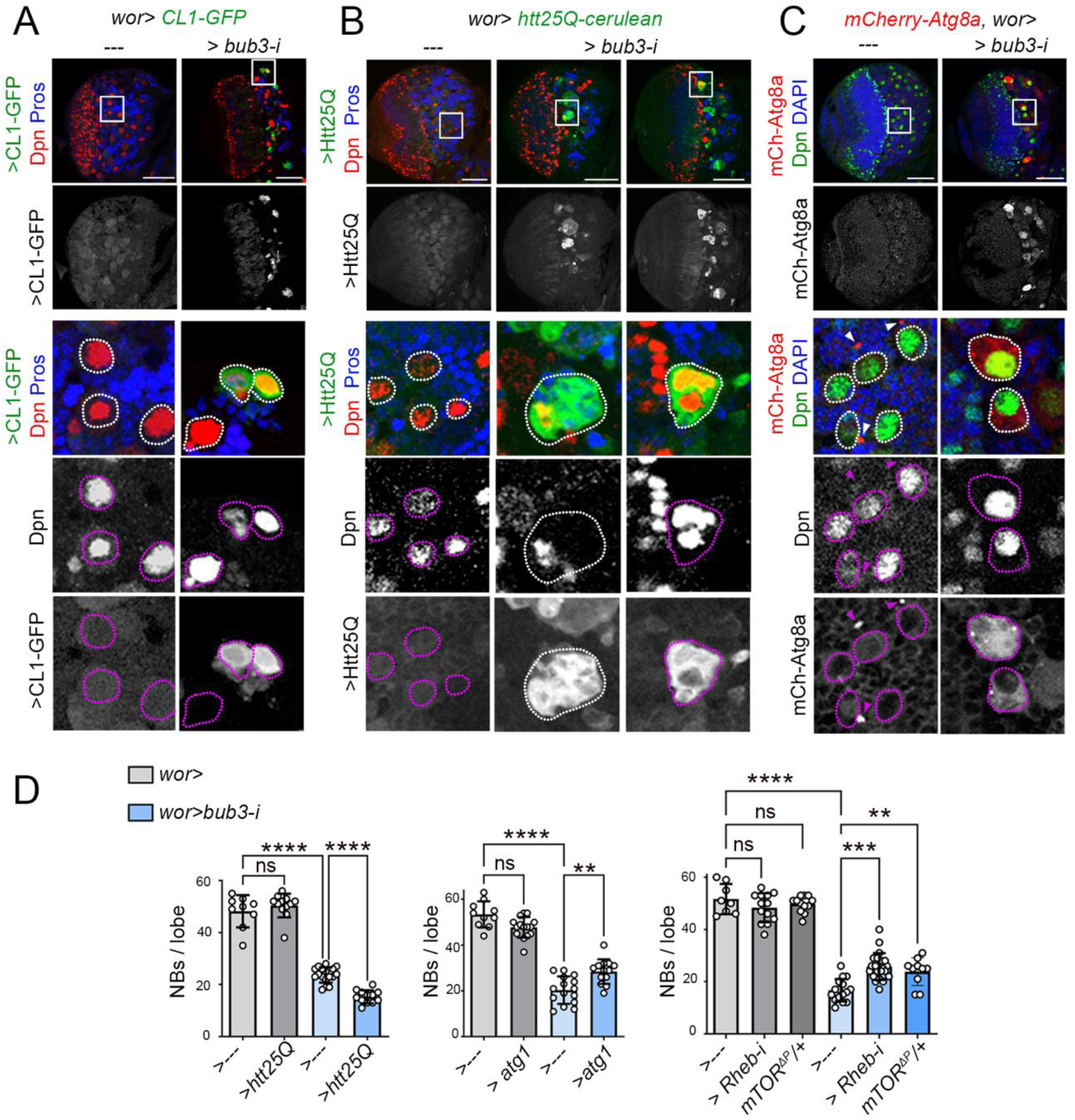
Proteostasis failure contributes to the loss of neural stem cells carrying complex aneuploidies. (**A**-**F**) Larval brain lobes expressing the indicated transgenes under the control of the *worniu-gal4* driver and labeled to visualize the expression of Dpn (red, green or white), Pros (blue, **A**, **B**), DAPI (blue, **C**), CL1-GFP (green or white, **A**), Htt25Q (green or white, **B**), and mCh-Atg8A (red or white, **C**). High magnifications of the squared regions are shown in the lower panels where NBs are marked by a dashed line. Scale bars, 50 µm. (**D**) Histograms plotting the number of NBs per brain lobe of larvae subjected to the expression of the indicated transgenes or heterozygous for *mTOR*. Mean and SD are shown. One-way ANOVA (**D**, Tukey’s multiple comparisons test) was performed:* p<0.05, ** p<0.01, *** p<0.001, **** p<0.0001 n.s., not significant. See also Figure S5 and Table S4.

The transcriptional upregulation of genes involved in mitochondrial disassembly and the transcriptional downregulation of genes encoding mitochondrial ribosome components (Figure 5C), together with the clear reduction in mitochondrial transcription (Figure 5B), pointed to a clear negative impact of aneuploidy on mitochondrial number or function. The developing brain is characterized by NBs strongly enriched by mitochondria surrounding the centrally localized nucleus (Figure 7A, B). In aneuploid NBs, we observed large accumulations of mitochondria that were not equally distributed around the nucleus (Figure 7A, B). Using a MitoTimer reporter, which targets the oxidation-sensitive Timer protein to mitochondria (Laker et al. 2014), we detected a clear shift of mitochondrial fluorescence toward red in aneuploid NBs (Figure 5C), which points either to high rates of mitochondrial oxidation or slow rates of mitochondrial turnover. We then used MitoQC (Lee et al. 2018), where a tandem GFP-mCherry fusion protein is targeted to the outer mitochondrial membrane, to analyze mitophagy, an important quality control mechanism that removes damaged mitochondria, in CIN brains. MitoQC exploits the pH sensitivity of GFP to enable differential labeling of mitochondria in the acidic microenvironment of the lysosome as a proxy endpoint readout. Only a few small MitoQC-positive puncta, most of them red, marking mitolysosomes, were observed in small cells around the NBs of control brains (Figure 7D, arrowheads). In contrast, large MitoQC-positive accumulations were observed in aneuploid NBs, and most (if not all) were positive for both colors (Figure 7D). We support the proposal that near saturation of autophagy compromises mitophagic activity, thus leading to the accumulation of dysfunctional mitochondria in aneuploid NBs. Consistent with this, overexpression of mitochondria-specific chaperones Hsp60 and Hsp60c, or ROS scavengers Sod2 and GTPx-1 was able to significantly rescue the number of aneuploid NBs observed in mature CIN brains, without causing any major effect in otherwise control brains (Figure 7E and Figure S6A). In the case of Hsp60 and Hsp60c, the number of Prospero-positive cells was also rescued (Figure 7E). These results indicate that mitochondrial dysfunction, most probably caused by near saturation of autophagy-compromising mitophagic activity, functionally contributes to the reduction of the NB population in CIN brains.

**Figure 7.**
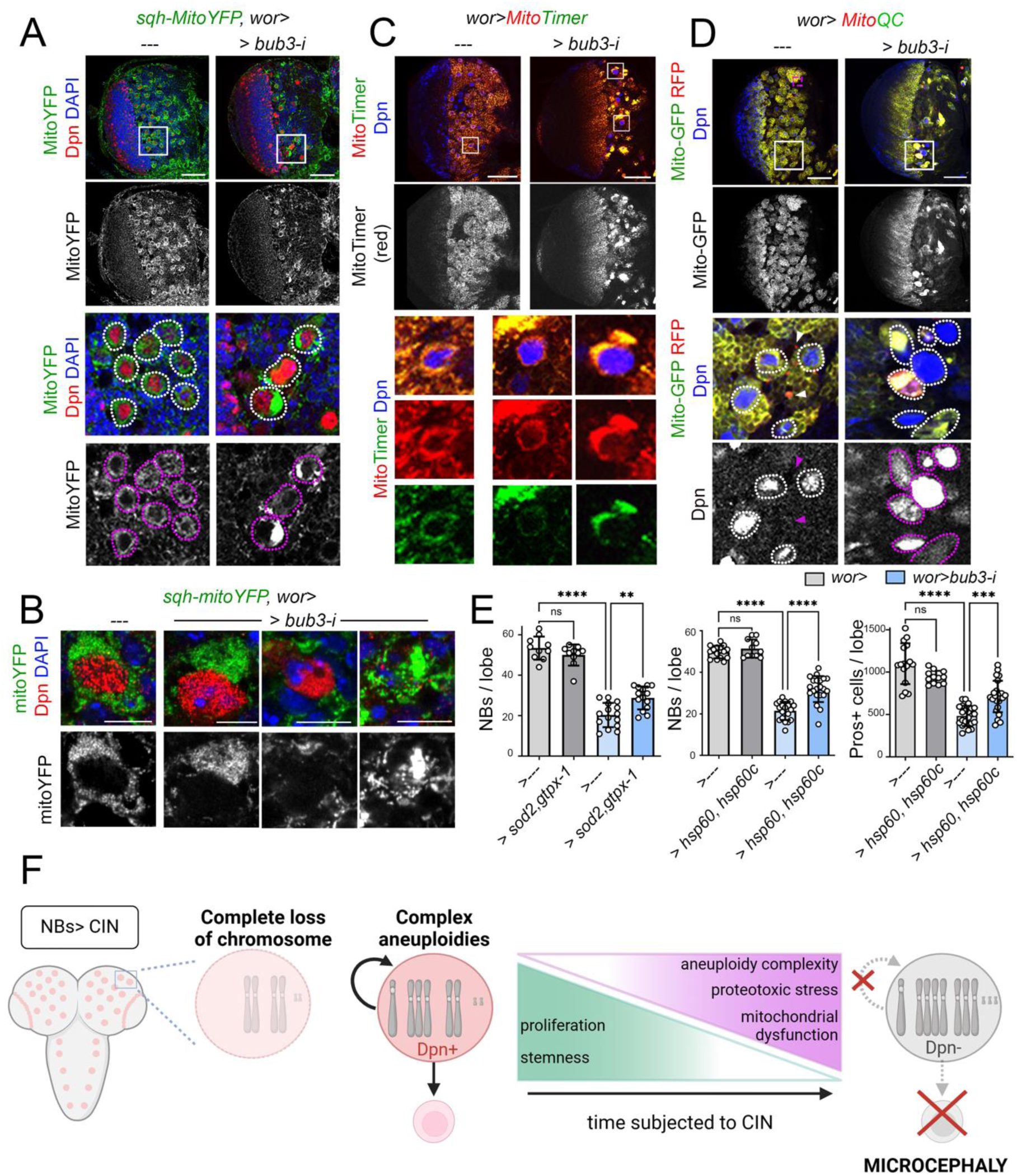
Mitochondrial dysfunction contributes to the loss of neural stem cells carrying complex aneuploidies. (**A**-**D**) Larval brain lobes expressing the indicated transgenes under the control of the *worniu-gal4* driver and labeled to visualize the expression of Dpn (red, blue and white), sq-Mito-YFP (green or white, **A**, **B**), Mito-Timer (green or white, **C**), and Mito-QC (green, red and white, **D**). In **A**, **C**, and **D**, high magnifications of the squared regions are shown in the lower panels and NBs are marked by a dashed line. Scale bars, 50 µm (A, C, D) and 7 µm (B). (**E**) Histograms plotting the number of NBs or Prospero-positive cells per brain lobe of larvae subjected to the expression of the indicated transgenes. Mean and SD are shown. One-way ANOVA (**E**, Tukey’s multiple comparisons test) was performed: * p<0.05, ** p<0.01, *** p<0.001, **** p<0.0001, n.s., not significant. (**F**) Cartoon summarizing the major cellular mechanisms underlying CIN induced microcephaly. Upon CIN induction in NBs, the accumulation of an unbalanced number of chromosomes consisting of multiple gains and/or losses leads to stemness loss (loss of NB identity and proliferative capacity) as a result of proteotoxic stress and mitochondria dysfunction. The complete loss of chromosomes is known to be cell lethal. See also Figure S6 and Table S4.

## Discussion

Mosaic variegated aneuploidy (MVA), a rare human disease that consists of widespread mosaic aneuploidies, is characterized by severe microcephaly (García-Castillo et al. 2008; Suijkerbuijk et al. 2010; Pavone et al. 2022; Malumbres and Villarroya-Beltri 2024). MVAs is caused by single mutations in genes involved in accurate segregation of chromosomes during mitosis (Matsuura et al. 2006; Hanks et al. 2004; de Wolf et al. 2021; Langeh et al. 2023; Yost et al. 2017; Carvalhal et al. 2022). Here we generated a *Drosophila* model of MVA-driven microcephaly by targeting the SAC genes *bub3* or *rod* specifically in the neural stem cell (NSC) compartment. We present evidence that loss of stemness—reduced viability and proliferative capacity—of SAC-depleted NSCs is the underlying cause of MVA-driven microcephaly and that it results in a lower number of neurons and glial cells. Using this model, we present evidence that it is the accumulation of an unbalanced number of gains and losses of more than one chromosome over time rather than a direct consequence of DNA damage or simple aneuploidies what compromises the identity and proliferative capacity of NSCs. This conclusion is consistent with previous reports unraveling a delayed response of NSCs to acute depletion of the cohesin complex, also known to cause chromosomal instability (Mirkovic et al. 2019). Thus, neither DNA damage caused by acute ionizing radiation had any impact on NSC identity or proliferative capacity, nor did the depletion of genes involved in the DNA damage response pathway strongly enhance the deleterious effects of SAC depletion on brain development. Similarly, despite the growth retardation observed in the developing brains of flies trisomic for the X and 2L chromosomes (carrying more than 2000 genes each), the number and proliferative activity of their NSCs were largely unaffected. Although highly aneuploid NSCs tended to express pro-differentiation factors, probably as a result of their identity loss, and some even activated the apoptotic pathway, their loss was neither a direct consequence of cell death nor a result of premature differentiation, as previously proposed (Gogendeau et al. 2015; Poulton, Cuningham, and Peifer 2017). Indeed, our transcriptomic analysis unveiled a strong effect of CIN on cellular homeostasis. Thus, the major responses of developing brains to CIN-induced aneuploidy included a reduction in genes involved in ribosome biogenesis and RNA metabolism, activation of protein quality control mechanisms (autophagy), and repression of mitochondrial activity. These transcriptomic changes are consistent with the conserved impact of aneuploidy on proteostasis (Stingele et al. 2012; Torres et al. 2007) and mitochondrial health (Joy et al. 2021). Thus, in yeast and human cells, aneuploidy leads to the production of an unbalanced proteome, which causes proteotoxic stress and activation of the major quality control mechanisms, including autophagy. Near saturation of autophagy in aneuploid cells has been demonstrated to compromise mitophagy, thus leading to the accumulation of dysfunctional mitochondria, which produce radical oxygen species (ROS). Consistent with the observed transcriptomic changes, highly aneuploid NSCs showed reduced expression of dMyc [a major regulator of ribosome biogenesis in vertebrates and invertebrates, (Pierce et al. 2008; Popay et al. 2021)] and smaller nucleoli, near saturation of the ubiquitin-proteasome system, strong activation of autophagy, and high sensitivity to protein aggregates. Activation of autophagy in these cells either by overexpression of Atg1 or depletion of TOR was able to partially rescue the reduced number of NSCs observed in SAC-depleted brains. Mitophagy was largely compromised in highly aneuploid NSCs, which also showed large accumulations of mitochondria not equally distributed around the nucleus. Overexpression of mitochondrial chaperones or ROS scavengers also partially restored the aneuploid NB count observed in mature CIN brains. We noticed that the number of aneuploid NBs was not fully rescued by any of these genetic interventions, most probably as a result of the highly pleiotropic effects of aneuploidy on cell physiology and on the fact that the complete loss of chromosomes is incompatible with cell viability. Nevertheless, those genetic interventions capable of dampening the deleterious effects of aneuploidy on NSC biology will undoubtedly offer new opportunities for drug discovery and disease treatment. Taken together, our data indicate that CIN-induced microcephaly stems, at least in part, from the generation, over time and through consecutive cell cycles, of NSCs carrying highly complex aneuploid karyotypes. Complex aneuploidies compromise the highly energy-demanding stemness of NSCs – their stem cell identity and proliferative capacity- by causing proteostasis failure and mitochondrial dysfunction (Figure 7F).

## Acknowledgments

We thank E. Baehrecke, N. Perrimon, J. B. Skeath, M. Suzanne, A. Whitworth, the Bloomington *Drosophila* Stock Center (USA), the Vienna Drosophila Resource Center (Austria), Kyoto Drosophila Stock Center (Japan), and the Developmental Studies Hybridoma Bank (USA) for flies and antibodies, Jery Joy for discussion, and IRB Barcelona’s Advanced Digital Microscopy, Functional Genomics and Biostastics/Bioinformatics Facilities for help. This work was funded by the PID2019-110082GB-I00 and PID2022-137673NB-I00 grants from the Spanish Ministry of Science and Innovation, and the ERDF “Una manera de hacer Europa”. We gratefully acknowledge institutional funding from the Spanish Ministry of Economy, Industry and Competitiveness (MINECO) through the Centres of Excellence Severo Ochoa Award, and from the CERCA Programme of the Catalan Government.

## Author Contributions

All authors conceived and designed the experiments and analyzed the data, AG-B, AAH and DB performed the experiments, M. M. supervised the whole project, analyzed the data, and wrote the paper.

## Declaration of interests

The authors declare no competing interests.

## Materials and Methods

### Key Resources Table

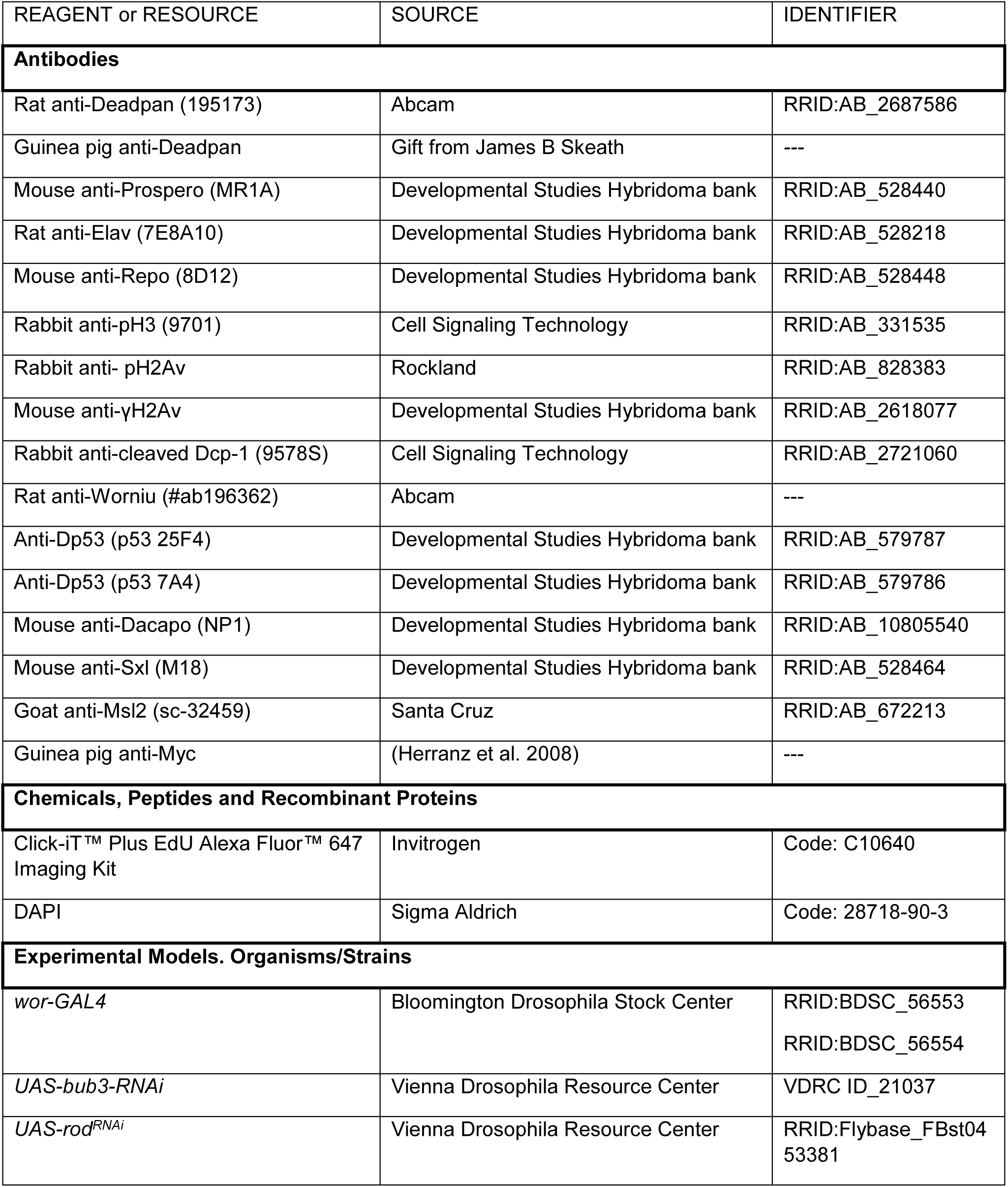
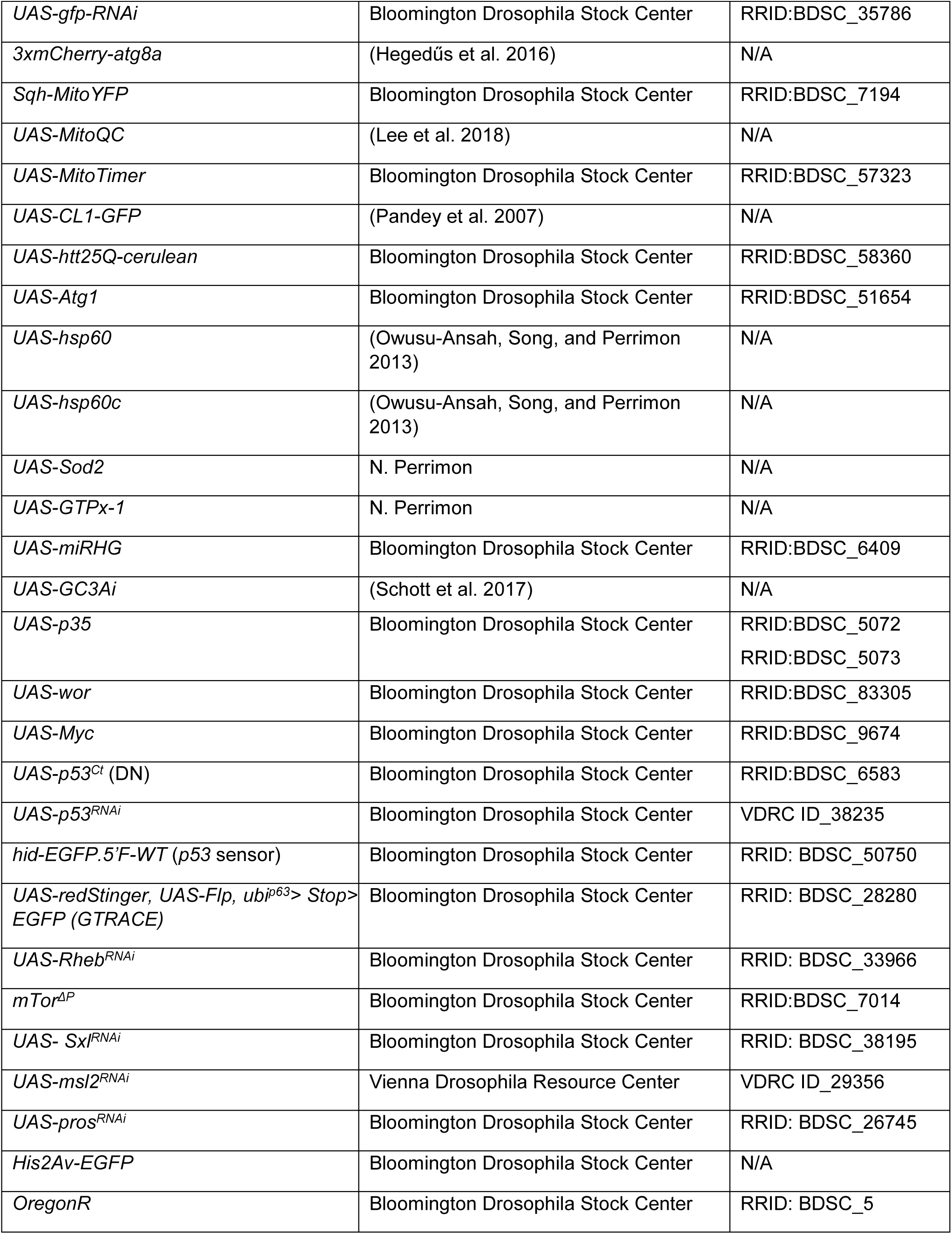
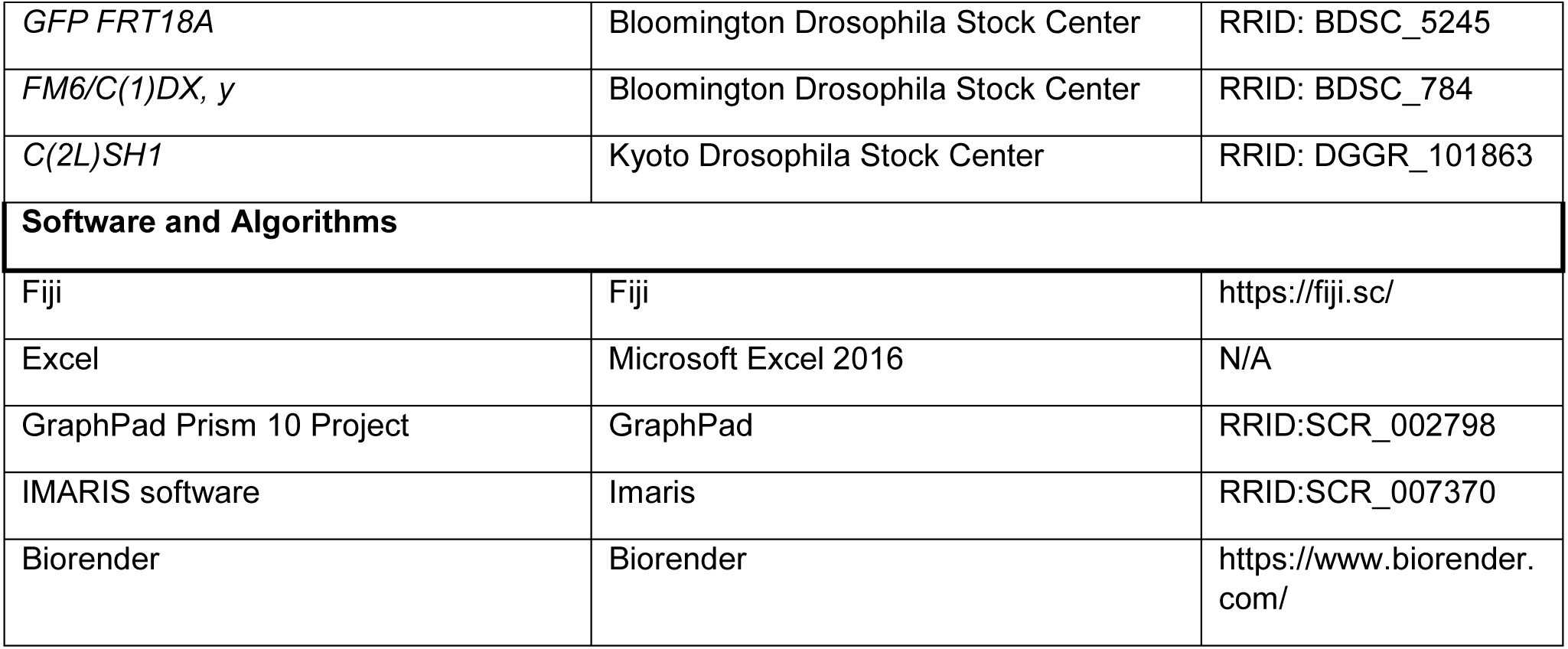

### Resource availability

#### Lead Contact

Further information and requests for resources and reagents should be directed to and will be fulfilled by the Lead Contact, Marco Milán.

#### Materials Availability

The strains generated in the course of this work are freely available to academic researchers through the Lead Contact.

### Experimental Model and Subject Details

#### Fly maintenance, husbandry and transgene expression

Strains of *Drosophila melanogaster* were maintained on standard medium (4% glucose, 55 g/L yeast, 0.65% agar, 28 g/L wheat flour, 4 ml/L propionic acid and 1.1 g/L nipagin) at 25°C in light/dark cycles of 12 h. When not specified, the sex of experimental larvae was not considered relevant to this study and was not determined. The strains used in this study are summarized in the Key Resources Table.

#### Standard induction of CIN and gene expression

Virgin female flies carrying the *wor-GAL4* driver and *UAS-bub3-i* were crossed with male flies carrying the gene of interest for each experiment. Fertilized females were allowed to lay eggs at 25^°^C and the resulting larvae were switched to 29°C 3 h after egg laying (AEL) in time point experiments, and 8 h AEL for the rest of the experiments. Brains were dissected after 48, 62, 72 or 96 h AEL. Both female and male larvae were used, except when characterizing Sxl and Msl2, in which females and males were considered separately. *UAS-gfp-RNAi* was used as a control for those experimental UAS transgenes targeting the corresponding pathways or genes.

#### Irradiation

Whole wild-type larvae were X-irradiated with 4500 rads in an YXLON MaxiShot X-ray system. Larval brains and wing discs were dissected and analyzed at the indicated time points after the irradiation.

#### Chromosome labeling by FISH

Fluorescence in situ DNA hybridization (FISH) was carried out to label the 2nd and 3rd chromosomes in brain cells. Briefly, larval brains were dissected in cold PBS and fixed with 4% paraformaldehyde for 20 min. Fixed tissues were washed with PBS-Tr (PBS1X; 0.3% Triton X-100) and transferred gradually into pre-Hybridization Mixture (pHM) (50% formamide; 4X SSC; 100mM NaH2PO4 pH7; 0.1% Tween 20). 150 ng of AACAC(2nd) -Alexa 488 -(AACAC)_6_) (Joyce et al. 2012) and dodeca (3rd)-Alexa 488-CCCGTACTGGTCCCGTACTGGT (Dekanty et al. 2012) chromosome probes were added and probe/chromosomal DNA was denatured by heating the sample to 91°C for 5 min. Hybridization was performed at 37°C overnight. After post-hybridization washes, brains were blocked and incubated overnight at 4°C in PBS-Tr; 10% Normal Goat Serum (NGS) with the primary antibody (anti-DPN). DAPI was used to counterstain DNA. Confocal sections covering the entire nuclei were taken and analyzed using Fiji Software (NIH, USA).

#### Immunohistochemistry

Larvae of interest were selected, and brains were dissected in phosphate-buffered saline (PBS), fixed for 20 min in 4% formaldehyde in PBS, and stained with antibodies in PBS with 0.3% BSA and 0.2% Triton X-100. Primary and secondary antibodies are summarized in the Key Resources Table.

#### DNA synthesis

Click-iT™ Plus EdU Alexa Fluor™ 647 Imaging Kit from Invitrogen (C10640) was used to measure DNA synthesis (S phase) in proliferating cells, following the manufacturer’s indications. EdU (5-ethynyl-2’-deoxyuridine) provided in the kit is a nucleoside analog of thymidine and is incorporated into DNA during active DNA synthesis. Larval brains were incubated with EdU for 20 min and then permeabilized in PBT of 40 min. The samples were incubated with the Click-iT reaction cocktails for 45 min at room temperature.

#### Generation of trisomic larvae

To generate trisomic larvae for the X chromosome*, C(1)DX, y* virgin females were crossed with *GFP-FRT18A* males (see Key Resources Table). Trisomic X female larvae were identified based on the absence of testes and the presence of GFP. To generate trisomic larvae for the 2L chromosome, *C(2L)SH1* virgin females were crossed with Oregon R males (see Key Resources Table). This cross produces both monosomic and trisomic embryos for the 2L chromosome. However, monosomic embryos do not survive past embryogenesis, ensuring that all larvae in the progeny are trisomic and available for dissection.

### RNAseq Data Processing and Availability

#### RNAseq library preparation

The samples used for this study were late third instar (L3) larvae from *Drosophila* (96 h AEL at 29 °C). The genotype of control larvae was *w; worniu-gal4, uas-gfp-RNAi*, while the genotype of the experimental larvae was *w; worniu-gal4, uas-bub3-RNAi*. Three independent biological replicates were performed on different days, with three brains dissected per genotype and replicate. All larvae used in this experiment were female. The brains were collected in 45 µl of lysis buffer containing 20 mM DTT, 10 mM Tris-HCl pH 7.4, 0.5% SDS, and 0.5µg/µl proteinase K, incubated at 65 °C for 15 min, and immediately frozen until processing. RNA isolation and library preparation were done at the IRB Barcelona Functional Genomics Core Facility. RNA was treated with DNAse I and purified using magnetic beads (RNAClean XP, Beckman Coulter). RNA quantity was measured with the Qubit RNA HS Assay kit (Invitrogen), and integrity was assessed with the Bioanalyzer 2100 RNA Pico assay (Agilent). Poly-A mRNA was purified from 150 ng of total RNA using the kit NEBNext Poly(A) mRNA Magnetic Isolation Module (New England Biolabs). Dual-indexed cDNA libraries were generated using the NEBNext Ultra II RNA Library Prep Kit for Illumina (New England Biolabs). All libraries underwent 13 min of RNA fragmentation and 13 cycles of PCR amplification. The final libraries were quantified using the Qubit dsDNA HS assay (Invitrogen) and subjected to quality control using the Tapestation HS D5000 assay (Agilent). An average size of 380 bp was confirmed. An equimolar pool was prepared with all the libraries for SE50 sequencing on a NextSeq 2000 (Illumina), and at least 26 million reads were obtained for each library.

#### Data processing

Adapters from reads were removed using trim galore (v.0.6.7) (https://github.com/FelixKrueger/TrimGalore) with default parameters. Resulting reads were aligned to the dm6 *D. melanogaster* genome version using STAR (v.2.7.10a) (Dobin et al. 2013). The count matrix was generated in R (https://www.R-project.org/) through the function “featureCounts” of the Rsubread package (Liao, Smyth, and Shi 2019).

#### Differential expression

Differential expression was performed using the DESeq2 package (Love, Huber, and Anders 2014). For plotting purposes and generating principal components, the count matrix was normalized using the “vst” function.

#### Gene set enrichment analysis

Gene set enrichment analysis was performed using the roastgsa package (Caballé-Mestres, Berenguer-Llergo, and Stephan-Otto Attolini 2023). GO and KEGG pathways were downloaded using the org.Dm.eg.db package.

#### Data availability

The raw and processed RNAseq data have been deposited in the GEO database under accession number GSE290206.

Reviewers can access the data at https://www.ncbi.nlm.nih.gov/geo/query/acc.cgi?acc=GSE290206 by entering the token ***odclwiiajdabbut*** in the access box.

### Image acquisition, Quantification and Statistical Analysis

#### Confocal microscopy

Samples were analyzed using the following confocal microscopes: LSM 780 Zeiss and LSM880 Zeiss fitted with a Fast Airyscan module (Carl Zeiss). Z-stacks were acquired using either the laser scanning confocal mode or the High-Resolution mode (Airyscan). Image acquisition was done with 40x and 63x oil lenses. The most representative image is shown in all experiments.

#### Brain size and cell quantification

Number of NBs was quantified manually using Fiji Software (NIH, USA) according to Dpn staining and nuclear size (>7µm). Only the NBs located in the ventral part of the central brain were taken into consideration. Size of the whole brain lobe, represented as area, was measured manually using Fiji Software on Z stacks obtained using a Zeiss LSM780 confocal microscope at 40x oil immersion objective. The numbers of progeny cells (GMCs, neurons and glia) were determined using IMARIS Software with the ‘Spots’ tool, according to Pros, Elav or Repo staining, respectively. Areas of G-TRACE and Ph2Av were measured with IMARIS software using the ‘Surface’ tool. *UAS-gfp-RNAi* was used as a control for those experimental UAS transgenes targeting the corresponding pathways or genes.

#### Statistical analysis

Statistical analysis was performed by unpaired equal-variance two-tail Student’s t-test when comparing the difference in means between two groups. We used a one-way ANOVA to compare three or more groups within the same experiment. In contrast, a two-way ANOVA was employed when there were multiple comparisons across different groups at various time points. Results for pairwise comparisons were adjusted by multiple contrasts using Tukey/Šídák correction, depending on the experiment (details in figures and Table S4). Differences were considered significant when p values were less than 0.0001 (****), 0.001 (***), 0.01 (**), or 0.05 (*). All genotypes included in each histogram or scatter plot were subjected to the same experimental conditions (temperature and time of transgene induction) and analyzed in parallel. Mean values and standard deviations were calculated and the corresponding statistical analysis and graphical representations were carried out with GraphPad Prism 10 statistical software.

## Notes

### Competing Interest Statement

The authors have declared no competing interest.

