## Supplemental Figures and Tables for "A pivotal contribution of proteostasis failure and mitochondrial dysfunction to chromosomal instability-induced microcephaly"

Amanda González-Blanco^1^, Adrián Acuña-Higaki^1^, David Boettger^1^, and Marco Milán^1,2,3 *^

**Supplementary Figures and Tables**

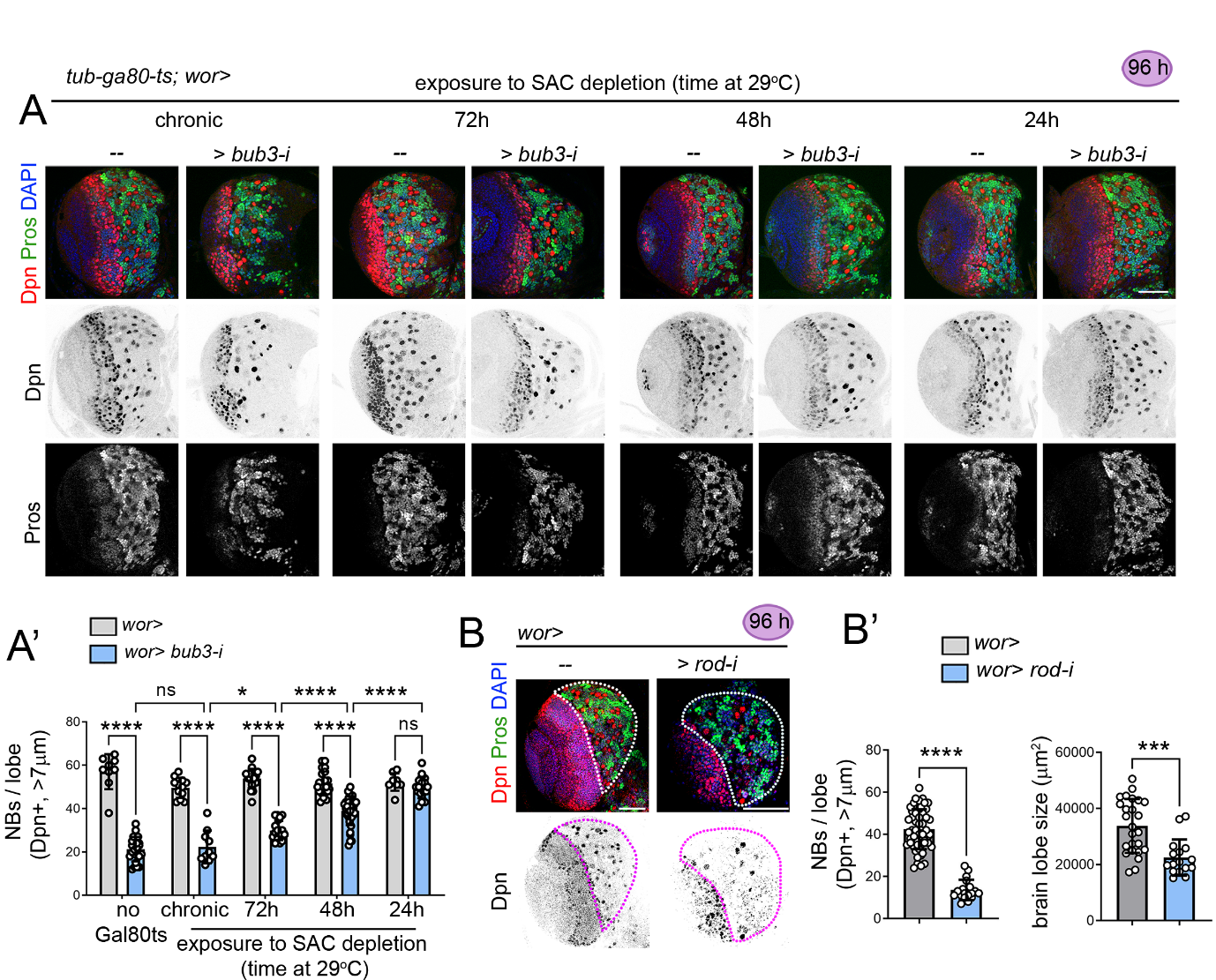

**Figure S1. Delayed response of neural stem cells to SAC depletion** (related to Figure 1).

(**A, B**) Larval brain lobes exposed to the expression of an RNAi form for *bub3* (**A**) or *rod* (**B**) under the control of the *worniu-gal4* driver, either chronically (since the embryo, **A**, first panels, and **B**), or during the indicated periods (in hours) and monitored at 96 h AEL to visualize the expression of Dpn (red and black), Pros (green and white) and DAPI (blue). Scale bars, 50 µm. (**A’**, **B’**). Histograms plotting the number of NBs per brain lobe, and the size of the brain lobe of larvae expressing the indicated transgenes and exposed to SAC depletion during the indicated periods (in hours). Mean and SD are shown. Two-way ANOVA (Šídák's multiple comparisons test) (**A’**) or Student’s t-test (**B’**) were performed: * p<0.05, ** p<0.01, *** p<0.001, ****p <0.0001, n.s., not significant.

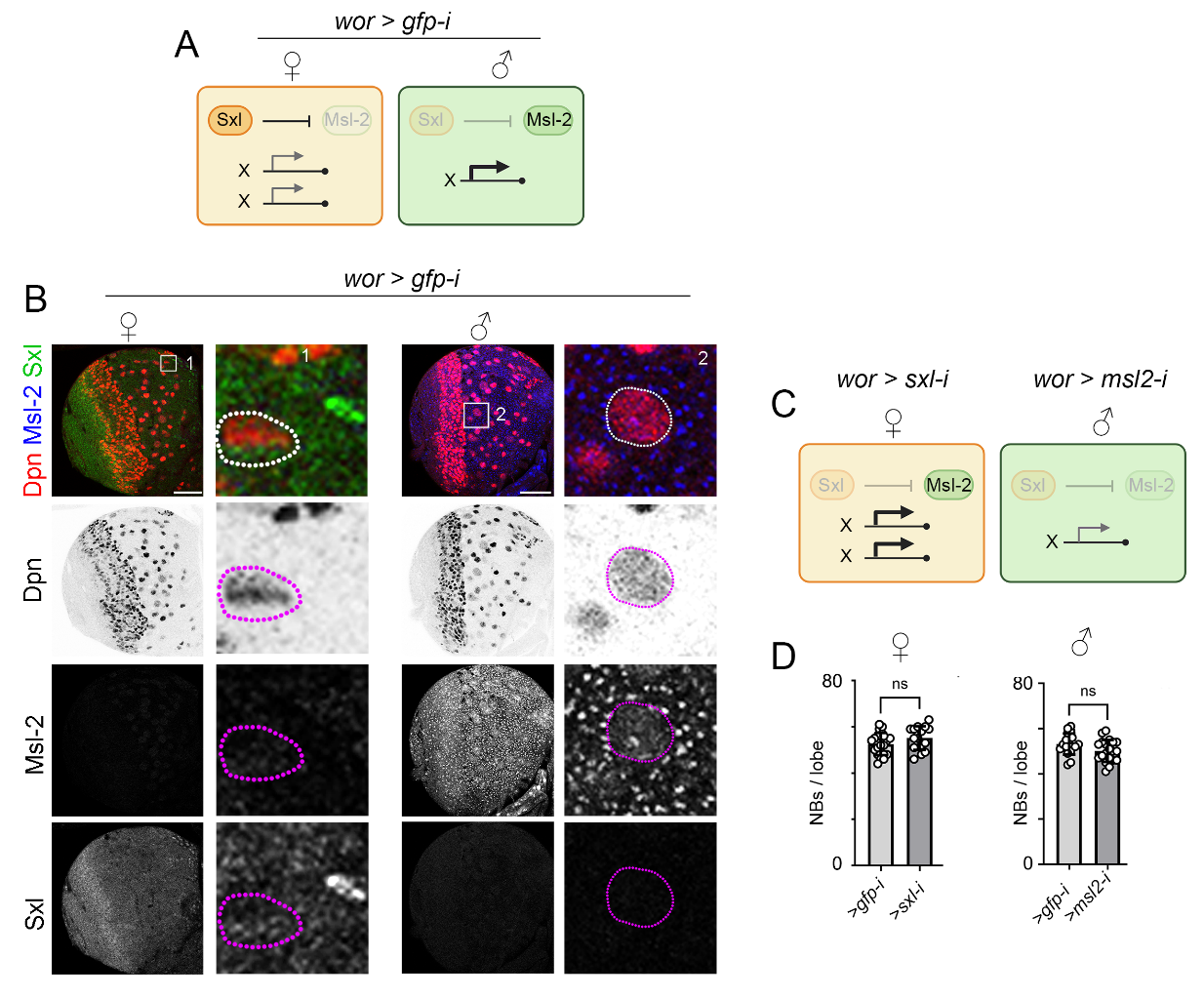

**Figure S2. Effects of gene dosage imbalance on neural stem cell viability** (related to Figure 3)**.**

(**A**, **C**) Cartoons depicting the expected expression levels of X-linked genes according to the presence of Sxl or Msl2 in the indicated sexes and genotypes. (**B**) Larval brain lobes of female and male larvae expressing the indicated transgene under the control of the *worniu-gal4* driver and labeled to visualize the expression of Dpn (red or black), Msl-2 (blue or white) and Sxl (green or white). NBs are marked by a dashed line in the high magnification of the squared regions. Scale bars, 50 µm. (**D**) Histograms plotting the number of NBs per brain lobe of female and male larvae expressing the indicated genotypes under the control of the *worniu-gal4* driver. Mean and SD are shown. Student’s t-test (**D**) was performed: * p<0.05, ** p<0.01, *** p<0.001, ****p <0.0001, n.s., not significant.

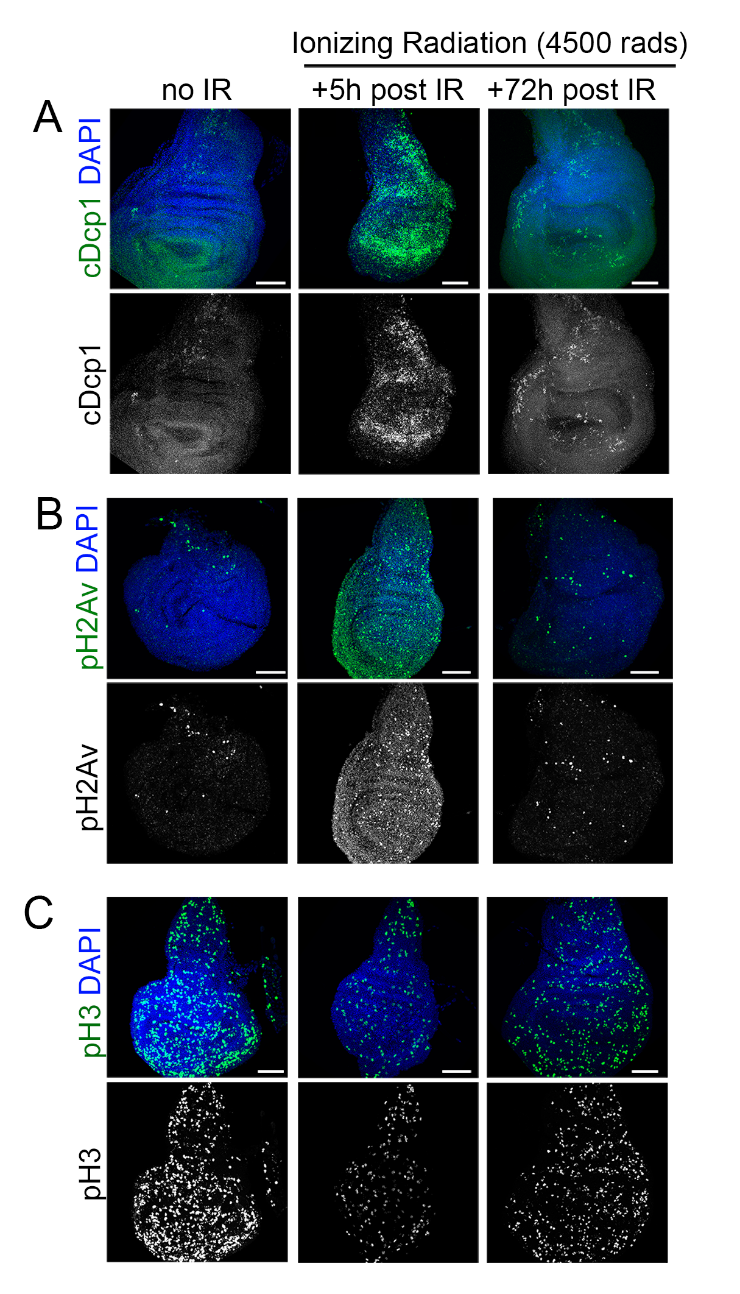

**Figure S3. Effects of Ionizing Radiation on larval epithelial cells** (related to Figure 4)

(**A**-**C**) Late third instar wing discs subjected to ionizing radiation 5 and 72 h before dissection and labeled to visualize DAPI (blue), cDcp1 (green and white, **A**), pH2Av (green and white, **B**), and pH3 (green and white, **C**). Untreated wing discs are shown in the first column. Scale bars, 50 µm.

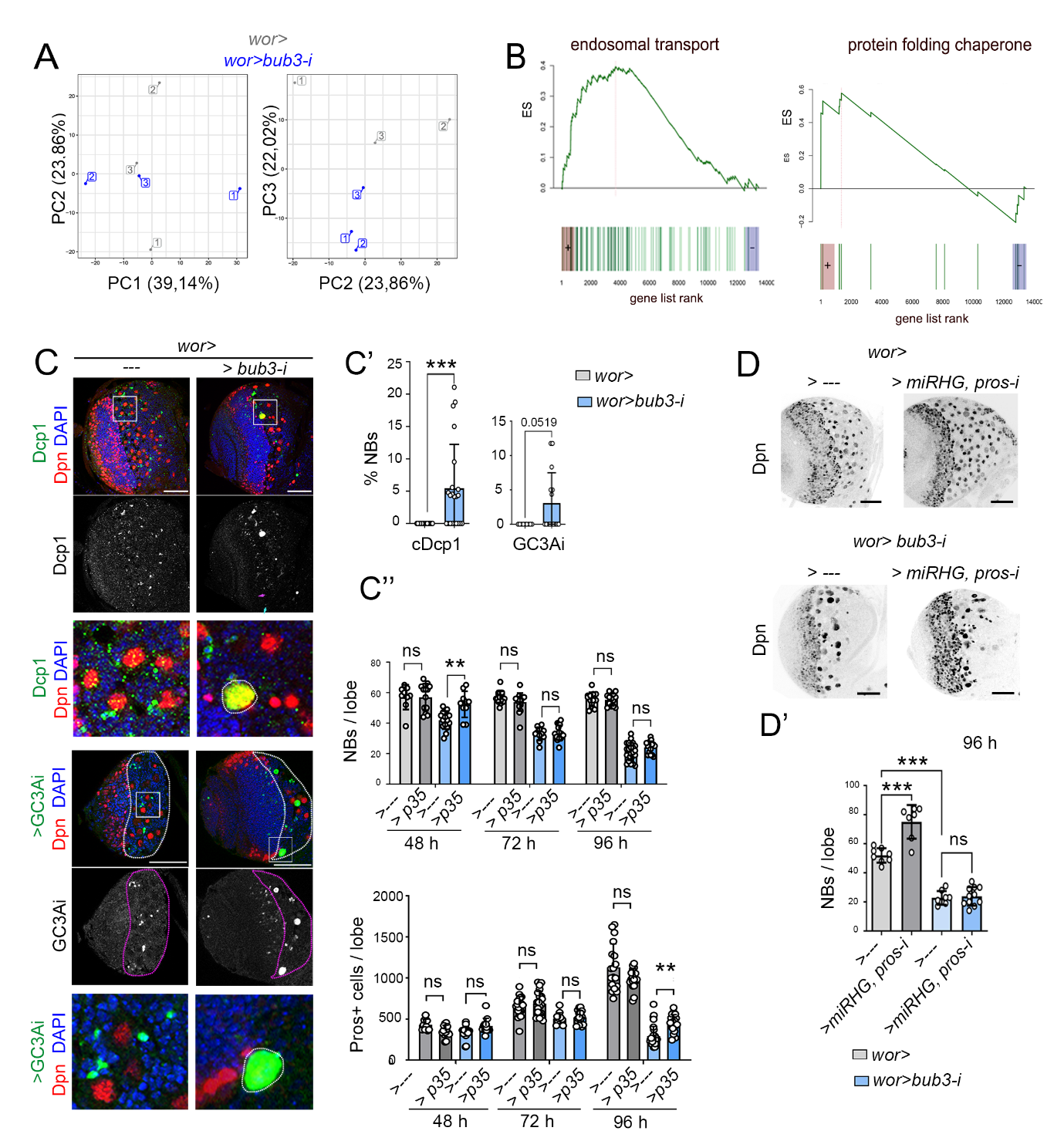

**Figure S4. Apoptosis-independent elimination of neural stem cells carrying complex aneuploidies** (related to Figure 5).

(**A, B**) Transcriptional changes of larval brain lobes exposed to chronic expression of an RNAi form of *bub3* (SAC depletion) when compared to controls (expressing *gfp-RNAi*) under the control of the *worniu-gal4* driver and monitored at 96 h AEL. In (**A**), Principal Components (PC) of the normalized expression matrix 1, 2 and 3 are shown. In (**B**), mountain plots of gene set enrichment analysis of significantly up- or down-regulated GO and KEGG pathways are shown. (**C**) Larval brain lobes expressing the indicated transgenes under the control of the *worniu-gal4* driver and labeled to visualize the expression of Dpn (red or black), cDcp1 (green and white, **A**, top panels), GC3Ai (green and white, bottom panels), and DAPI (blue). In **C**, high magnifications of the squared regions are shown in the lower panels and NBs are marked by a dashed line. Scale bars, 50 µm. **(C’)** Histograms plotting the percentage of NBs labeled with cDcp1 or GC3Ai, (**C’’**) the number of NBs and Prospero-positive cells per brain lobe of larvae subjected to the expression of the indicated transgenes in time points. Mean and SD are shown. Mann Whitney test (C’), two-way ANOVA (C’’, Tukey’s multiple comparisons test), and one-way ANOVA (D’, Tukey’s multiple comparisons test) were performed: * p<0.05, ** p<0.01, *** p<0.001, ****p< 0.0001, n.s., not significant.

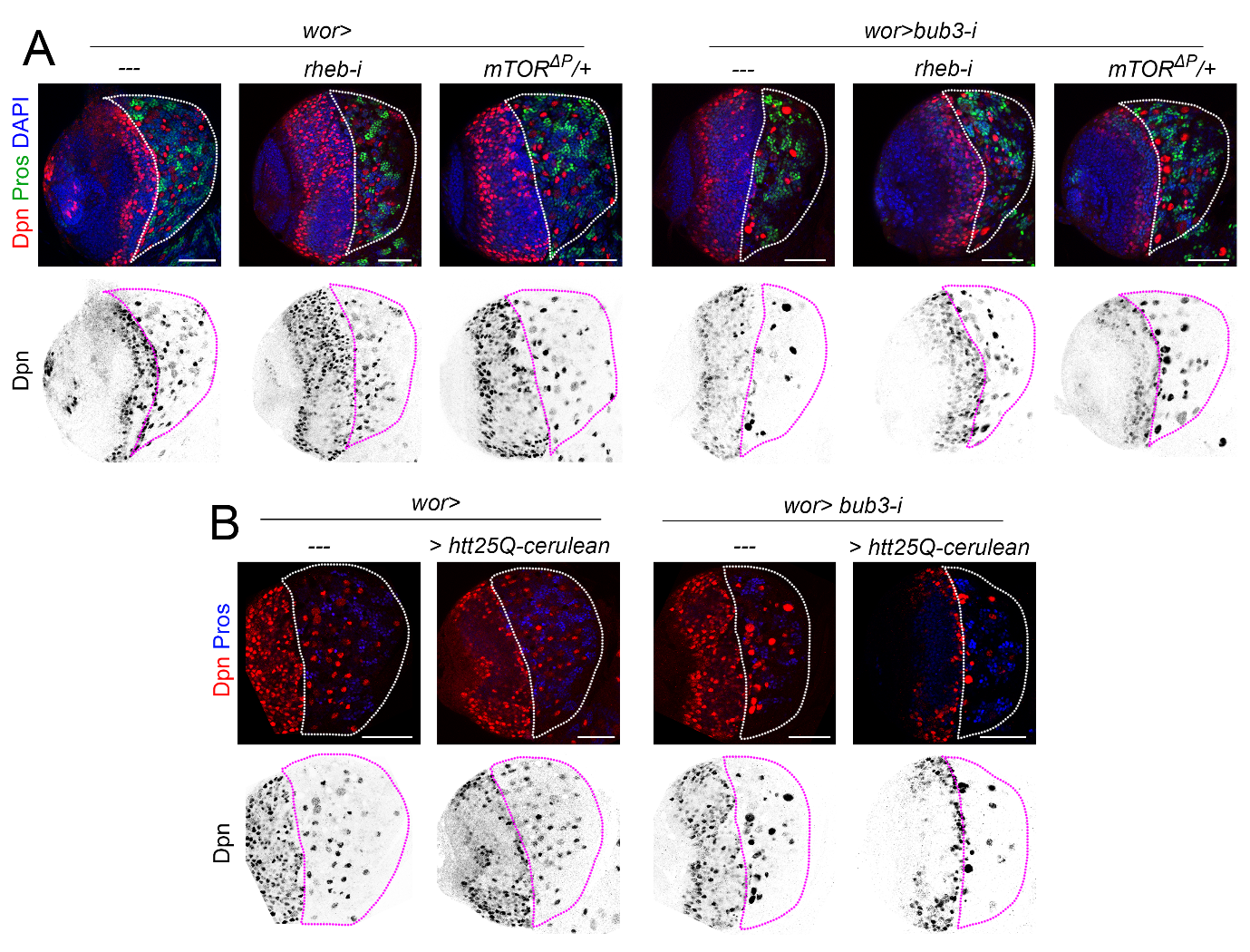

**Figure S5. TOR depletion rescues the loss of neural stem cells carrying complex aneuploidies** (related to Figure 6)

(**A**, **B**) Larval brain lobes expressing the indicated transgenes under the control of the *worniu-gal4* driver and labeled to visualize the expression of Dpn (red and black), Pros (green), and DAPI (blue). Scale bars, 50 µm.

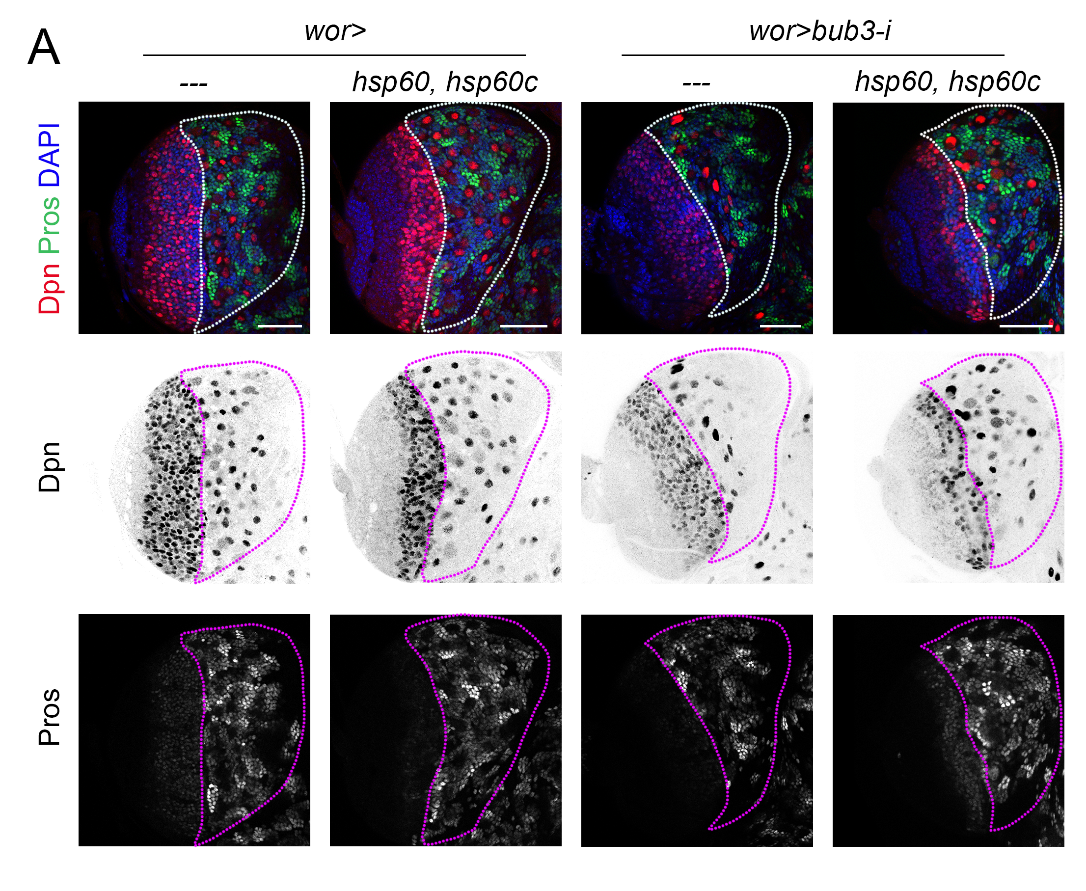

**Figure S6. Mitochondrial chaperones rescue the loss of neural stem cells carrying complex aneuploidies** (related to Figure 7)

(**A**) Larval brain lobes expressing the indicated transgenes under the control of the *worniu-gal4* driver and labeled to visualize the expression of Dpn (red and black), Pros (green), and DAPI (blue). Scale bars, 50 µm.

**Table S1. Transcriptional changes of larval brains exposed to CIN: single genes** (related to Figure 5). List of genes significantly up- or down-regulated in larval brains subjected to CIN when compared to controls.

**Table S2. Transcriptional changes of larval brains exposed to CIN: GO and KEGG pathways** (related to Figure 5). List of GO and KEGG pathways significantly up- or down-regulated in larval brains subjected to CIN when compared to controls.

**Table S3. Transcriptional changes of larval brains exposed to CIN: genes within GO and KEGG pathways** (related to Figure 5). List of genes of each of the GO and KEGG pathways significantly up- or down-regulated in larval brains subjected to CIN when compared to controls.

**Table S4. File containing the parameters quantified and the statistical details** (related to Figures 1-7 and S1-S6). **Quantification and statistical analysis of all measurements performed.**
